# Combinatory differentiation of human induced pluripotent stem cells generates thymic epithelium that supports thymic crosstalk and directs dendritic- and CD4/CD8 T-cell full development

**DOI:** 10.1101/2023.12.23.572664

**Authors:** Nathan Provin, Manon d’Arco, Erwan Kervagoret, Alexandre Bruneau, Lucas Brusselle, Cynthia Fourgeux, Jérémie Poschmann, Xavier Saulquin, Carole Guillonneau, Laurent David, Matthieu Giraud

**Affiliations:** Nantes Université, CHU Nantes, Inserm, Centre de Recherche Translationnelle en Transplantation et Immunologie, UMR 1064, F-44000 Nantes, France; Nantes Université, Inserm, CNRS, Centre de Recherche en Cancérologie et Immunologie Intégrée Nantes Angers, 44035 Nantes, France

**Keywords:** Thymus, iPSc, differentiation, thymic organoids, thymopoiesis

## Abstract

The thymus is a primary lymphoid organ playing a crucial role in immune tolerance, by educating thymocytes through a selection mediated by thymic epithelial cells (TEC). Recent advances in gene editing and immunotherapies have made in vitro generation of iPSC-derived T cells a crucial issue with promising therapeutic applications. Current state-of-the-art approaches often fail to accurately replicate the thymic niche, resulting in impaired T cell generation, particularly for CD4+ T cells. Here we address the production of functional mature iPSc-derived TECs that are able to support in vitro T cell generation. We designed a protocol allowing iPSc differentiation into thymic epithelial progenitors (TEP) through an unbiased multifactorial method based on optimal experimental design. Modulation of signaling pathways known to regulate embryonic thymus organogenesis resulted in the obtention of TEPs expressing typical thymic markers. We achieved TEP maturation into medullary and cortical TECs by setting up a hydrogel-based 3D culture supplemented in RANK ligand, resulting in the formation of thymic organoids. To assess the functionality of the generated TECs, primary hematopoietic progenitors were co-cultured in the organoids and showed significantly improved maturation in single-positive CD4 or CD8 T cells. Remarkably, our thymic organoid model shows multilineage differentiation potential, with generation of distinct dendritic cells populations in addition to the T lineages. Thus, generation of functional TECs and thymic organoids offers a practical platform for the study of thymic cellular interactions and paves the road to future cellular therapies by producing mature thymic immune cells of clinical interest.

## Introduction

The thymus plays a crucial role in the establishment of central immune self-tolerance. Its main function is the generation of a diverse yet non-autoreactive T lymphocyte repertoire (Miller, 2020). The thymus is subdivided in several lobes structured in two principal niches, a peripheral cortical area surrounding a central medulla. Distinct cellular populations shape specific microenvironment of these niches. Thymic epithelial cells (TECs) are a crucial actor that regulates the maturation of developing T cells, the thymocytes (Abramson and Anderson, 2016). Two main populations of TECs are identified: the cortical TECs (cTECs) and the medullary TECs (mTECs) that are characterized in human by the KRT8/18 and KRT5/14 cytokeratin expression. Although KRT8/18 tag all epithelial cells, KRT14 is restricted to the medullary subset and KRT5 to most mature TECs. In addition, the expression of the surface marker CD205 marks cTECs (Abramson and Anderson, 2016; Kadouri et al., 2019). Recently, single-cell studies have revealed a higher level of heterogeneity for TECs with, notably, the identification of new mTEC subpopulations with various functional roles such as thymocyte homing, commitment, proliferation regulation and selection (Bautista et al., 2021; Bornstein et al., 2018; Kadouri et al., 2019). Thymocyte selection is required for the generation of a functional and non-autoreactive T repertoire and is mediated in two steps. Thymocytes originate from a CD34+ circulating hematopoietic early progenitor (ETP) and differentiate through several stages characterized by CD4 and CD8 expression (Carpenter and Bosselut, 2010; Chopp et al., 2020; Deng et al., 2021). Positive selection is mediated by cTEC at the double positive (DP) stage and leads to the selection of thymocytes with a functional T cell receptor (TCR) (Kadouri et al., 2019). Negative selection filters out potential autoreactive thymocytes at the single-positive (SP) stage in the medulla (Klein et al., 2014, 2000) and is mediated by HLA-DR^hi^ mTECs expressing a great diversity of peripheral tissues antigens (PTAs) under the regulation of AIRE (Anderson et al., 2002; Cumano et al., 2019; Kadouri et al., 2019). Mature CD4 and CD8 T lymphocytes are called recent thymic emigrants and migrate to the periphery (Cosway et al., 2021). Recent research has shown that parallel intrathymic dendritic cell (DC) differentiation was occurring from the same ETPs (Liang et al., 2023). Interestingly, crosstalk with this thymic DC population has been shown to support back thymocyte maturation and selection (Liang et al., 2023; Spidale et al., 2014).

mTECs and cTECs derive from a common bipotent progenitor with cortical-like phenotype (Alves et al., 2014; Baik et al., 2013). Molecular mechanisms regulating TEC fate decision are still elusive, although crucial implication of NOTCH (Li et al., 2020; Liu et al., 2020) and RANK-CD40-LTB signaling (Irla, 2021; Irla et al., 2008; Lopes et al., 2022), provided by thymocytes through the thymic crosstalk, have been reported. Regulation of the organogenesis of the thymus has also known remarkable recent progress, notably thanks to the publication of single cell RNA sequencing (scRNA-seq) thymic cell atlas at different stages of development (Gao et al., 2022; Park et al., 2020; Zeng et al., 2019). Thymic epithelial progenitors (TEP) identity is characterized by expression of *FOXN1*, the master regulator inducing expression of the TEC transcriptional program (Blackburn et al., 1996; Bredenkamp et al., 2014; Nowell et al., 2011). TEPs emerge from the third pharyngeal pouch endoderm (3PPE), a transient embryonic structure formed by the involution of the endoderm inside the foregut tube between week 3 and 4 in human (Farley et al., 2013; Gordon et al., 2001; Gordon and Manley, 2011). A gene cascade involving *TBX1*, *PAX9* and *EYA1* has been identified as a crucial regulator of 3PPE formation. 3PPE is derived from the ventral pharyngeal endoderm, itself originating from the anterior foregut endoderm (AFE) (Farley et al., 2013; Figueiredo et al., 2020). Regulation of the fate acquisition in these structures has also shown recent progress at the single-cell level (Han et al., 2020; Magaletta et al., 2022; Nowotschin et al., 2019).

Mimicking thymus organogenesis *in vitro* to generate TECs from embryonic (ESc) and induced pluripotent stem cells (iPSc) has been a long-standing objective over the last decade, using both reprogramming and mainly direct differentiation approach mainly (Bredenkamp et al., 2014; Chhatta et al., 2021; Gras-Peña et al., 2022; Parent et al., 2013; Ramos et al., 2022; Soh et al., 2014; Sun et al., 2013). However, the complexity of the regulation of the differentiation process and our limited comprehension of *in vivo* mechanisms hindered the achievement of this goal (Provin and Giraud, 2022). Indeed, thymic differentiation often yields immature and non-functional cells with a progenitor (TEP) identity and requires grafting *in vivo* to complete the maturation (Parent et al., 2013; Ramos et al., 2022; Sun et al., 2013). Moreover, the available protocols of iPS differentiation into TEPs have not been optimized by unbiased systematic approaches to robustly assess the effects of the supplemented factors at each stage (Callaghan et al., 2022; Toms et al., 2017; Yasui et al., 2021). Thus, reliable methods for differentiation of iPSc into functional mature TECs *in vitro* and for their long-term maintenance in a culture system reproducing the thymic niche are yet to be developed. Such systems would result in the generation of iPSc-derived TECs, opening promising perspectives for regenerative medicine (Besnard et al., 2021).

*In vitro* T Lymphocyte generation has been greatly improved using feeder stromal cells such as OP9-DLL1 or MS5-hDLL4, modified to express NOTCH ligand (Bosticardo et al., 2020; Montel-Hagen et al., 2020; Seet et al., 2017). Formation of artificial thymic organoids improved the maturation state of obtained T cells, and partially reproduce thymic function (Chhatta et al., 2021; Iriguchi et al., 2021). However, these setups could only partially reproduce thymic selection, lacking the self-antigen presentation ability assured by mTEC^hi^.

Because of the complexity of thymic organoids co-cultures and iPSc thymic differentiation caused by the multiplicity of factors involved, precise optimization of the protocols is hard to achieve. Here we used optimal design of experiments (DOE), a systematic statistical framework for experimental design to collect the most information with the fewest experiments. Transcriptomics readout was used to optimize iPSc thymic differentiation based on recent single cell datasets of thymus organogenesis. We propose a two-week protocol that efficiently yields TEPs expressing thymic identity markers. To overcome the difficulty to mature TEPs *ex vivo*, we developed a 3D coculture system with primary thymocytes (ETPs). We set up a fibrin hydrogel air-liquid interface culture that resulted in the formation of human thymic organoids (hTOs). hTOs could be maintained up to 5 weeks, and showed evidence of TEC maturation, notably into HLA-DR^hi^ mTECs, with higher *AIRE* expression than cells cultured in monolayer. T-TEC physical interactions were demonstrated, and hTOs showed the ability to drive ETP maturation into SP CD4^+^ and CD8^+^ T lymphocytes. Further characterization of hTO products by scRNA-seq confirmed the mature stage of the generated T lymphocytes. Finally, we report secondary differentiation of ETPs into dendritic cell lineage in line with recent observations *in vivo,* thus demonstrating a complex *in vitro* organoid model with both cell maturation and multilineage potential.

## Materials and methods

### Isolation of human primary early thymocyte progenitors (ETP) and thymic epithelial cells (TEC)

Postnatal human thymic samples were obtained as anonymized, discarded waste from patients undergoing cardiac surgery at Maternité de Nantes, in accordance with the French CODECOH regulation under declaration DC-2017-2987. ETPs isolation was performed as described in Lavaert *et al*. (Lavaert et al., 2020). Briefly, fresh thymus samples were dissected on the same day in 1 mm^3^ fragments in RPMI1640 and dissociated by mechanical pipetting to release thymocytes. A 5-minute incubation in red blood cell lysis solution (Miltenyi, 130-094-183) was performed for erythrocyte depletion. Cells were passed through 70µm meshes and incubated with CD4 and CD8 labeled Dynabeads (Thermofisher, 11031) to deplete most DP and SP thymocytes. ETP-enriched cell suspension was stained for flow cytometry using antibodies referenced in supplementary methods. ETPs were sorted with a FACS ARIA (BD Biosciences) using phenotype CD3^-^CD4^-^CD8^-^CD14^-^CD19^-^CD56^-^CD34^+^CD7^+^ and immediately used for reaggregation culture. For TEC isolation, thymic fragments were digested using a 0.5 mg.mL^-1^ collagenaseD (Roche, 11088866001), 1 mg.mL^-1^ dispase (Roche, 04942078001) and 0.5 mg.mL^-1^ DNAseI (Roche, 11284932001) solution in RPMI1640 for 45 minutes at 37°C and dissociated using GentleMACS (Miltenyi, 130-093-235) and C tubes (Miltenyi, 130 093 237). Cell suspension was deposited on a 21% Optiprep (Sigma Aldrich, D1556 250ML) gradient and centrifuged 20 min at 500g for TEC enrichment. Cell suspension was filtered on a 100µm mesh and sorted with phenotype CD45^-^EPCAM^+^.

### iPSc culture and differentiation in thymic epithelial progenitors (TEP)

iPSc cell lines were furnished by the Nantes iPSc platform and were maintained on Matrigel (Stem Cell Technologies, 354277) coated 6-well plates in mTESR1 medium (Stem Cell Technologies). Details on the cell lines used are available in Supplementary figures. iPSc cultures were passaged every 5 to 6 days, using XF passaging solution (Stem Cell Technologies) and were seeded in clumps. In case of anormal colony morphology in cultures, daily cleaning was performed to guarantee optimal iPSc purity. To control cell state variability, iPSc were passaged 4 days before induction of differentiation, then harvested 24 hours before the D0 in single cell suspension using TrypLE (ThermoFisher) with 5 minute incubation at 37°C. Cells were resuspended in mTESR1 medium with 10µM ROCK inhibitor Y27632 (Sigma Aldrich Y0503) and seeded on Matrigel coated 12-well plates at 37.10^3^ cells/cm^2^. Differentiation was induced at D0 with XVIVO10 medium (Lonza, BE04-380Q) supplemented with 5µM CHIR99 (Miltenyi, 130-106-539) and 100 ng.mL^-1^ ActivinA (BiotechneR&D, 338-AC). Culture medium changes were performed at fixed hours, with quick rinsing by warm PBS to eliminate potential differentiation medium leftovers in wells. D1 and D2 medium contained only 50 ng.mL^-1^ ActivinA. At D3, cells are passaged in single cell as described above and seeded at low density (14.2 to 20.10^3^ cells/cm^2^ depending on iPSc line) in Matrigel coated 12-well plate with 10 µM Y27 and 50 ng.mL^-1^ ActivinA. Culture medium was changed at D4, D5, D7, D9, D10, D11 and D13 as previously described with supplementation as detailed in Figure 1.

**Figure 1.**
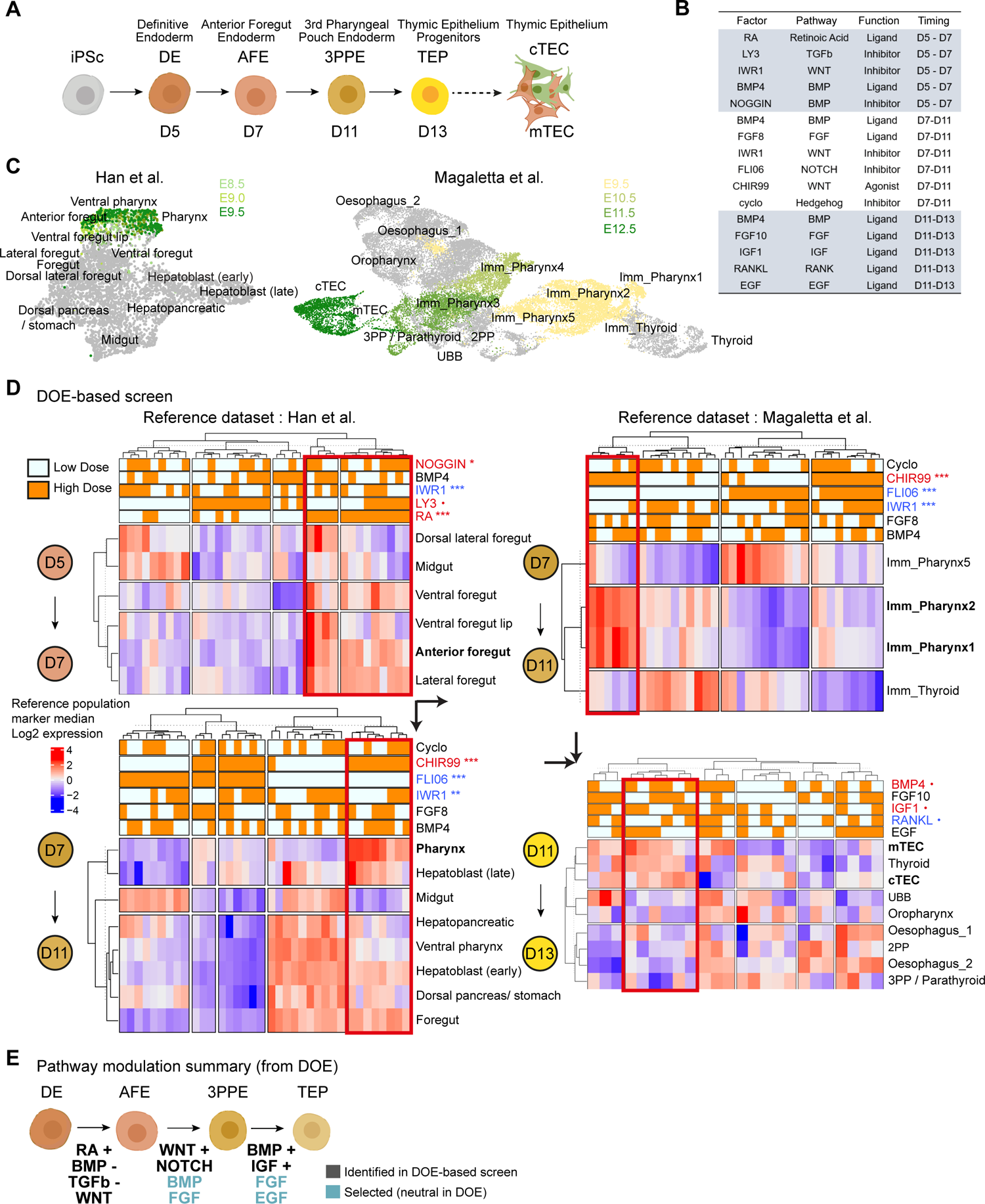
Combination of DOE and DEG-seq results in optimized thymic differentiation from iPSc. **(A)** Differentiation itinerary of the thymic epithelium. **(B)** Factor abbreviations, impacted pathways and timings for the experimental design screening. **(C)** UMAP representation of reference data with identify the transcriptomic signature of the thymus organogenesis differentiation trajectory (yellow to green)**. (D)** DGE-seq results for each DOE experiment on the expression score of target clusters in the differentiation trajectory. Statistical significance is measured by ANOVA. Significant factors were selected for the optimized protocol. **(E)** Synthesis of the pathways identified by the DOE analysis. Black: + activation, -inhibition, no sign indicates the pathways that are necessary and shouldn’t be inhibited. Blue: neutral DOE pathways but selected.

### TEP reaggregation with early thymic progenitors (ETP) and thymic organoids (hTO) formation

TEP were harvested at day 13 using TrypLE with a 7-minute incubation at 37°C and mechanical flushing. TEP were aggregated in low binding 96-well plates (ThermoFisher, 174929), at a 8:1 ratio with fresh ETP sorted as described above. Culture is maintained 24 hours in an XVIVO10-based medium supplemented with 1% Glutamax (Gibco, 35050-38/2165269), 1% non-essential amino acids (Gibco, 11140-035), 30 mM L-ascorbic () 0.1µM retinoic acid (Sigma Aldrich, 302-79-4), 50 ng.mL-1 BMP4 (Miltenyi, 130-111-165), RANKL (BiotechneR&D: 6449-TEC), 10 ng.ml-1 FGF8 (BiotechneR&D, 423-F8), FGF10 (Miltenyi, 130-127-858), IGF1 (Miltenyi, 130-093-886), EGF (Miltenyi, 130-097-751), SCF () and 5ng.mL-1 IL7 (), FTL3L (). Cell micromasses must show a typical spheroid morphology with a compact central cell mass surrounded by a crown composed mainly of ETPs. Hydrogels were formed by mixing a 8 mg.mL^-1^ Fibrinogen (Sigma Aldrich, 341578) solution heated at 37°C with 10 U.mL^-1^ Thrombin (Sigma Aldrich,605190) and 26000 U.mL^-1^ Aprotinin (Sigma Aldrich, 616370), and immediately casting it in the superior compartment of 24-well inserts. 2-4 spheroids were individually collected using cut 200µL pipette tips pre-coated with anti-adherence solution and seeded delicately on the hydrogels. The same culture medium described above was added in a way to form an air-liquid interface without drowning the hydrogels and changed every three days during the first week. Then retinoic acid, BMP4, RANKL, FGF8, FGF10, IGF1, and EGF were removed from the culture medium. To harvest the cells, the hydrogels were carefully collected using cut 1000-µL pipette tips and incubated 15 minutes at 37°C in a 0.5 mg.mL^-1^ Collagenase (Roche, 11088866001) solution in DMEM/F12. Organoids were subsequently processed with a 10-minute incubation in TrypLE at 37°C with frequent mechanical disruption by pipetting, then filtered on a 100µm mesh.

### RNA-extraction, RT and qPCR

Cell samples were lysed using RLT Lysis buffer (Qiagen, 79216). RNA extraction was performed using RNAeasy mini kit (Qiagen, 74104) and retrotranscription with Superscript III (Thermofisher, 18080093) following the manufacturer’s guidelines. qPCR was performed with SYBR Green Fast kit (ThermoFisher, 4385612) on a ViiA 7 Real-Time PCR System (ThermoFisher) using the primers listed in supplementary methods. Relative quantification of target gene expression was measured using the delta CT method with *GAPDH* as endogenous control.

### Bulk RNA sequencing by DGE-seq and SMART-seq

For 3’ Digital gene expression (DGE) bulk RNA sequencing, protocol was performed according to our implementation of previously developed protocols (Kilens 2018). 10 ng total RNA was used for library preparation. mRNA was tagged thanks to poly(A) tails-specific adapters, well-specific barcodes and universal molecular identifiers (UMI) with template-switching retrotranscription. cDNAs were pooled, amplified and tagmented by transposon-fragmentation to enrich 3’ ends. Sequencing was performed on Illumina® HiSeq 2500 using a Hiseq Rapid SBS Kit. Kits used were Zymo purification kit Researche D4004-1-L, Kit Advantage 2 PCR Enzyme System (Clontech, 639206), QIAquick Gel Extraction, Kit AgencourtAMPure XP magnetic beads, Nextera DNA (FC-121-1031). For the deep SMART-seq RNA sequencing, RNA quality was first controlled with TapeStation (Agilent D1000), then the processed with SMART-Seq v4 Ultra Low Input RNA kit, and the library prepared with Illumina Nextera XT DNA Library Preparation Kit. Illumina NextSeq500 was used for the paired-end sequencing (2 × 150 bp). Reads were homogenized to 2 × 100bp by read-trimming and mapped with TopHat2.

### RNA-seq data processing and analysis

Read pairs used for analysis matched the following criteria: all 16 bases of the first read had quality scores of at least 10 and the first 6 bases correspond exactly to a designed well-specific barcode. The second reads were aligned to RefSeq human mRNA sequences (hg19) using bwa version 0.7.17. Reads mapping to several transcripts of different genes or containing more than 3 mismatches with the reference sequences were filtered out from the analysis. DGE profiles were generated by counting for each sample the number of unique UMIs associated with each RefSeq genes. DGE-sequenced samples were acquired from five sequencing runs.

Sequenced samples with at least 50000 counts and 6000 expressed genes were retained for further analysis. Batch correction and differentially expression (DE) was performed with DEseq2 R package, with the threshold of 1 log2 fold change and 0.05 Benjamini Hochberg p-value to classify a gene as differentially expressed vs D0 iPSc controls. Gene Ontology of DE genes modules was performed using ClusterProfiler R package on KEGG database of biological processes. GO network was plotted using REVISEGO and Cytoscape software. Data visualization relied on custom pipelines based on ComplexHeatmap and ggplot2 R package.

### scRNA-seq data generation

D28 dissociated hTO were sorted on FACS ARIA to purify cell population and remove dead cells and debris. Three samples were labeled by HTO staining and processed with a Chromium Single-Cell 3’ Reagent v2 Kit (10x Genomics, Pleasanton, CA) as per the manufacturer’s protocol. Briefly, a small volume of the single-cell suspension was mixed with RT-PCR mix and loaded with partitioning oil and Single Cell 3’ Chip. Chip was loaded on Chromium controller (10X Genomics) for single cell generation and barcoding. cDNA was then pooled, pre-amplified and fragmented. Adapters were incorporated and sequenced with Illumina® HiSeq 2500.

### scRNA-seq data processing and analysis

Primary data analysis was performed on CellRanger. Cell count matrices were exported and secondary analysis was performed on R using Seurat v4 package (Hao et al., 2021). HTO were demultiplexed and quality control defined by keeping cells with gene number between 2000 and 15000 and fewer than 0.1 mitochondrial genes proportion. Variable gene identification, data scaling and dimension reduction used functions integrated in Seurat. Dimensionality reduction used UMAP and tSNE embedding on the first 10 dimensions. Clustering used the Louvain algorithm implemented in FindCluster function with a resolution of 0.3. Visualization and marker determination relied on default Seurat functions. For the analysis of public sc datasets of thymus and pharyngeal organogenesis, raw data was downloaded at the accession provided in the data availability section of the relevant papers (21,31,32). Between-sample integration was performed using the integration framework provided by Seurat. Data analysis followed the same pipeline. Park’s dataset was downsized by random sampling by a factor of 0.25 to ease data handling. Quality control was set at 500 / 10000 / 10, and 500 / 4000 / 10 for Han’s and Magaletta’s datasets. Dimension reduction was performed by UMAP on first 10 (Park), 21 (Magaletta) and 25 (Han) dimensions. Clustering was performed with a resolution of 0.75 (Park), 0.5 (Magaletta), 0.6 (Han). Visualization and marker identification were performed as described above. For label transfer of our dataset to the thymus atlas reference dataset, anchor identification, integration and transfer relied on the default Seurat vignette.

### Flow cytometry and Immunofluorescence (IF) imaging

Cells were stained with antibodies listed in supplementary according to the manufacturer’s guidelines and analyzed on FACS ARIA (BD Biosciences) and Celesta (BD Biosciences). For 2D IF, cells were cultivated on Ibidi 8-well plates (Ibidi, 80806), fixed with 4% PFA for 15 minutes and permeabilized with IF buffer (PBS, 0.2% Triton, 10% inactivated FBS) for 1 hour. Cells were incubated in IF buffer overnight at 4°C with primary antibody and for 2 hours at room temperature for secondary antibodies. Plates were imaged on a confocal SIM microscope. For hTO stainings, the hydrogels were harvested by using a widely cut 1000µL pipette tip pre-coated with anti-adherence solution and carefully deposited in Ibidi 8-well plates in cold PBS. Most of the gel matrix was removed by delicate flushing and several PBS washing, without damaging the main cellular structures. hTOs were fixed with 4% PFA for 20 minutes and permeabilized with IF buffer (PBS, 0.2% Triton, 10% inactivated FBS) for 2 hours. Primary staining was performed in IF buffer for 24h and secondary staining for 4 hours, and imaging on SIM confocal microscope in Z-stack mode.

### Statistics and data analysis

Statistical tests were performed on R. Mean comparison significance on qPCR graphs was tested by non-parametric t.tests with a significance p-value threshold of 0.05. DOE experimental plans were designed using R DOE base package 1.11.6. DOE results were analyzed by factorial ANOVA as detailed in figure legends.

## Results

### Optimization of directed iPSc differentiation towards a TEC fate using DOE-based factor screening and bulk transcriptomics

To set up an optimized differentiation protocol of iPSc into TEPs, we followed an approach of directed differentiation recapitulating the events of the thymic organogenesis. Recently, the understanding of thymic identity acquisition and its underlying regulation at the cellular and molecular level has greatly improved through the use of sc-OMIC approaches (Han et al., 2020; Li et al., 2021; Magaletta et al., 2022; Nowotschin et al., 2019). Definitive endoderm (DE) anteriorizes into anterior foregut endoderm (AFE) from which emerges the third pouch pharyngeal endoderm (3PPE), a transitory structure that gives rise to the thymic epithelial progenitors (TEPs) (**Figure 1A**). We aimed to recapitulate these events *in vitro* by modulating the main pathways involved in thymic organogenesis, such as BMP, WNT, Hedgehog (SHH) and FGF.

Exit of pluripotency state and DE induction have been thoroughly studied (D’Amour et al., 2005) and the process of iPSc differentiation into DE precisely characterized. We performed this step of differentiation by stimulating Nodal through ActivinA exposure in combination with a 24-hour pulse of WNT through CHIR99. The differentiating cells showed a peak of expression of the DE marker *SOX17* at days 3-5 (**Supp Figure 1A**). In addition, a vast majority of cells stained positive for SOX17 and another DE marker, FOXA2, at day 5 (**Supp Figure 1B**). These results reflect the high efficiency of the iPSc to DE differentiation that does not require further optimization.

We applied DOE to test the effect of a series of factors impacting differentiation pathways and potentially active in the transitions towards AFE, 3PP and TEP. The modulation of RA, TGFb, BMP and WNT pathways on definitive endoderm anteriorization between D5 and D7 was first studied. Then the effect FGF8, NOTCH, Hedgehog (SHH), BMP and WNT modulation of 3PPE was assessed between D7 and D11. Lastly, the effect of BMP, FGF10, EGF, IGF1 and RANKL modulation on TEP fate induction between D11 and D13 was investigated (**Figure 1B**). These pathways were selected based on studies of thymic organogenesis regulation (Balciunaite et al., 2002; Barbarulo et al., 2016; Bleul and Boehm, 2005; Davenport et al., 2016; Provin and Giraud, 2022) and previous thymic differentiation of hESc (Parent et al., 2013; Sun et al., 2013). A Plackett-Burman combinatorial screening design was performed for each of the three transitions by testing multiple combinations of factors with two modalities of concentration (**Supp Figure 1, D**).

Using a low-dimensionality readout, such as measuring the expression of one or a few marker genes, can yield imprecise results due to marker lack of specificity or gene expression variability. To overcome this weakness and robustly assess factor effects, we performed bulk RNA-seq on the DOE samples and compared their transcriptome to two scRNA-seq atlases of pharyngeal development. The one by Han et al. (Han et al., 2020) corresponds to early stages of development and covers the AFE and 3PPE transitions. The second by Magaletta et al. (Magaletta et al., 2022) focuses on later stages of development including the 3PPE and TEC transitions. Raw data reanalysis of these two datasets, in applying uniform manifold approximation and projection (UMAP), led to the delineation the transcriptomic maps of thymic differentiation trajectory from E8.5 to 9.5 (Han) and E9.5 to 12.5 (Magaletta) (**Figure 1C**) through the study of the pattern of expression of known genes specific to thymic precursor cell populations (**Supp Figure 2**). We then scored the transcriptome similarity of each DOE sample to relevant *in vivo* thymic precursor cell populations at each stage of the trajectory using mean expression of marker genes of each cell population, defined as the top 100 genes the most specifically expressed.

**Figure 2.**
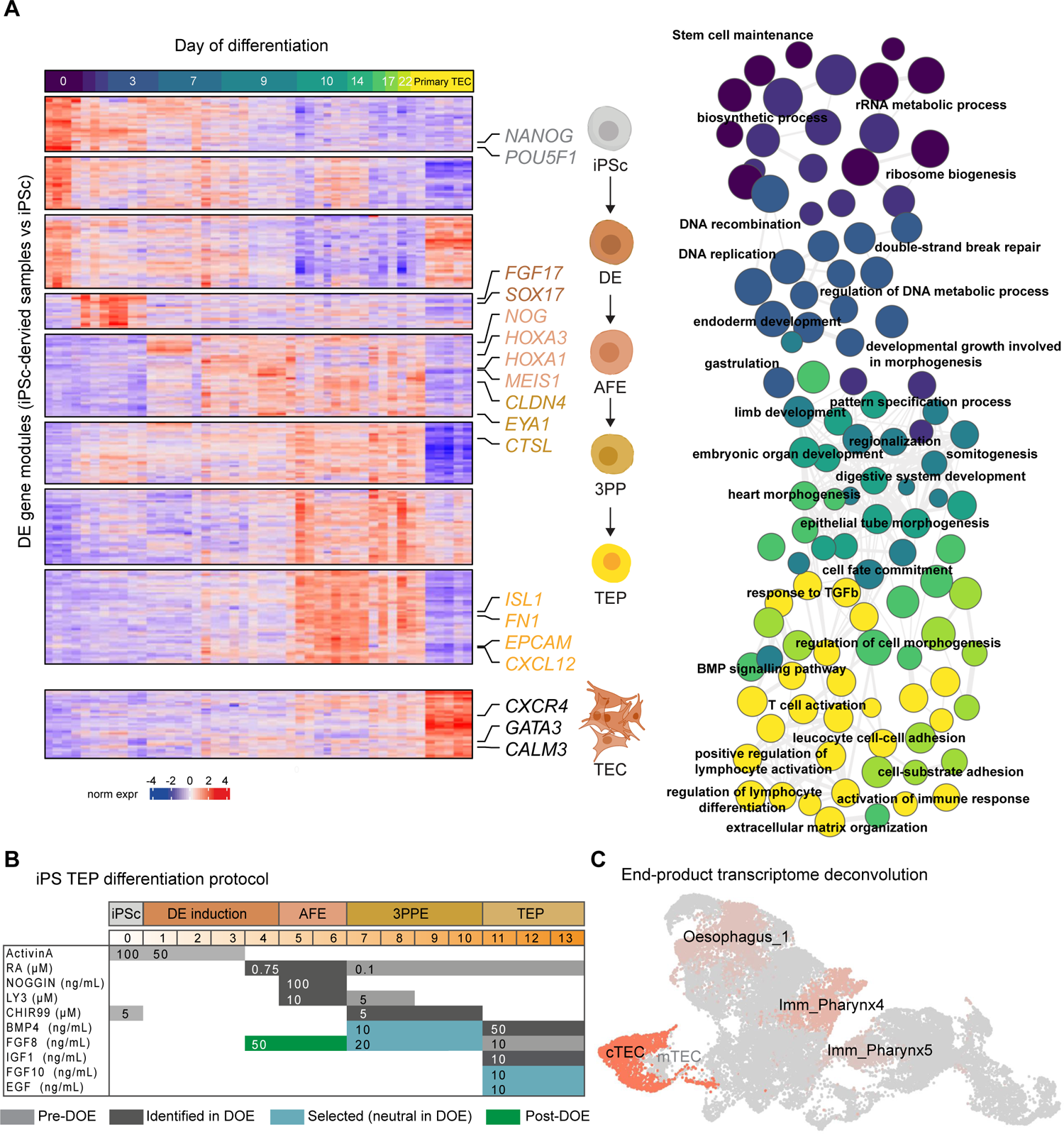
Transcriptomic characterization of iPSc-derived TEP. **(A)** Heatmap representation of differentially expressed genes modules across differentiation (*Left*). Sample were analyzed at several timepoints by DGE-seq. Primary human TECs and D0 iPSc were included as controls. GO terms enriched in each gene module were identified by ClusterProfiler and organized by shared genes similarity (*Right*). Network was plotted on Cytoscape by forcing orientation along the day of differentiation, resulting in the graph of significant biological processes enriched for each differentiation stage. **(B)** Recapitulation table of our optimized protocol. **(C)** Deconvolution of the end-product of differentiation onto the Magaletta reference dataset of pharyngeal organogenesis.

The differentiation conditions into AFE were optimized against Han dataset. Unsupervised hierarchical clustering of DOE samples and thymic progenitor cell populations identified two clusters of samples showing high expression for anterior foregut markers (**Figure 1D, Top Left**). These samples also show a strong expression of markers of the developmentally close populations: lateral foregut and ventral foregut, as well as low expression of midgut and dorsal lateral foregut markers, confirming DE anteriorization. To identify the effect of the tested factors on anterior foregut induction, we computed their statistical main effects by factorial ANOVA on the mean expression of the anterior foregut gene markers. RA, NOGGIN and LY3 supplementation, as well as absence of IWR1, showed the most impact on DE anteriorization (**Supp Figure 3**). Hence, BMP inhibition by NOGGIN and, to a lesser extent, TGFβ inhibition by LY3 promoted anteriorization, whereas WNT inhibition by IWR1 is globally detrimental at this step.

**Figure 3.**
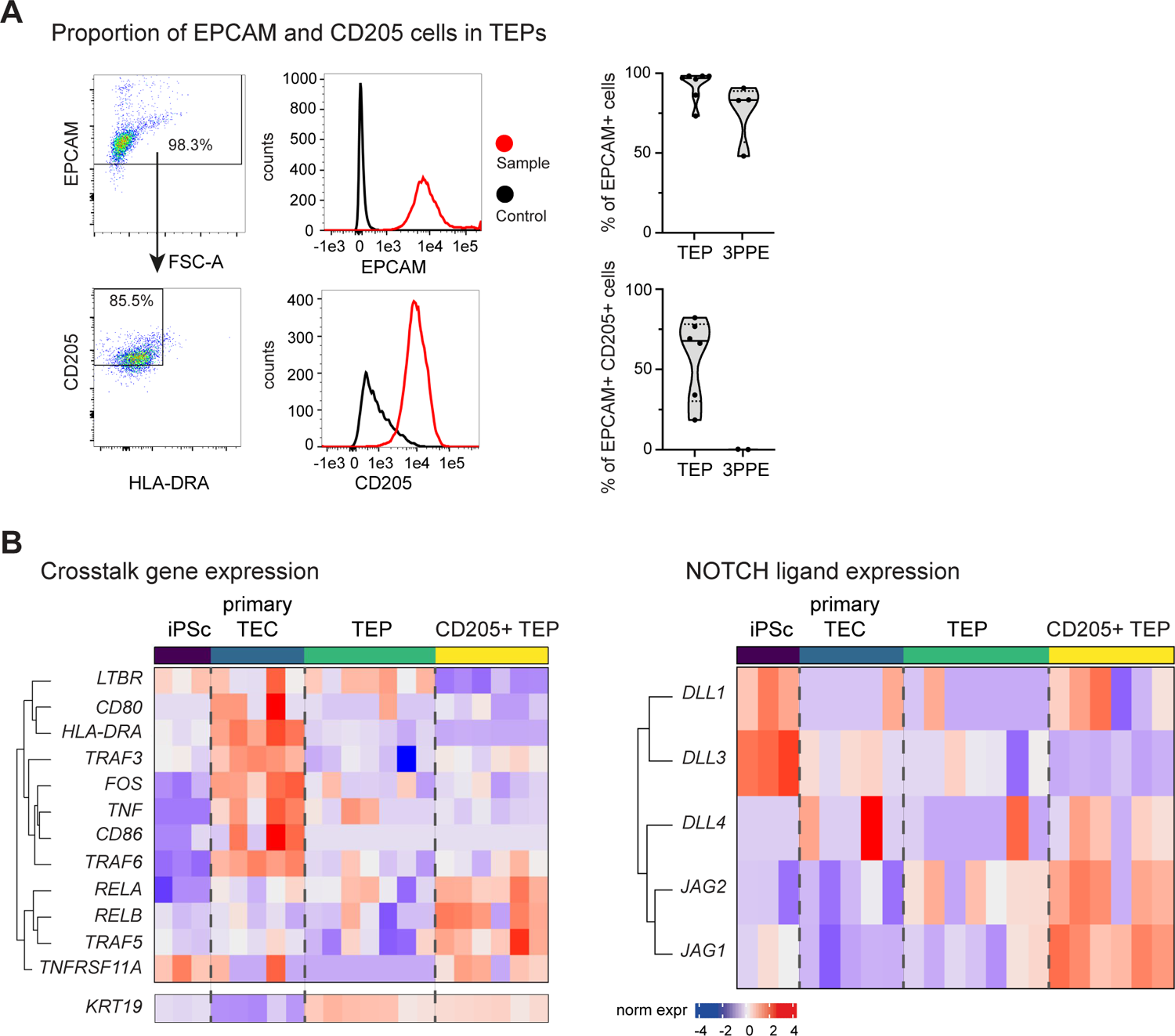
Identification of the potential of TEPs to receive and provide key developmental signals. **(A)** Quantification of TEPs expressing EPCAM and the TEP/cTEC lineage marker CD205. **(B)** Transcriptomic data of D14 -D22 TEP show expression of genes involved in the thymic crosstalk (*Left*) and NOTCH pathway activation in thymocytes, with DO iPSc and primary TECs as controls

We applied a similar strategy to the 3PPE transition and found a cluster of DOE samples with the highest expression for pharynx markers (**Figure 1D, Bottom Left**). The ANOVA of the tested factors on the mean expression of the pharynx gene markers identified a statistically significant effect for the addition of CHIR99, as well as the deprivation of FLI06 and IWR1 (**Supp Figure 3**). We then used the Magaletta dataset as reference, since it also encompasses the transition to 3PPE with immature pharynx 1 and 2 populations that correspond to the developmental stage of the pharynx population of Han dataset. We confirmed the positive effect of CHIR99, as well as the negative effect of FLI06 and IWR1 on 3PPE formation (**Figure 1D, Top Right; Supp Figure 3**). Hence, WNT activation by CHIR99, in contrast to its inhibition by IWR1, greatly promotes pharyngeal formation of the 3PPE, whereas NOTCH inhibition by FLI06 is detrimental. Interestingly, contrary to Parent *et al*. study (Parent et al., 2013), we observed no effect of Hedgehog inhibition by cyclopamine at this stage. Similarly, BMP4 and FGF8 supplementation showed no significant effect for pharyngeal induction. However, we chose to keep BMP4 and FGF8 because of their consensual use to achieve 3PPE differentiation in previous studies.

Finally, we optimized the differentiation conditions of the transition to TEP in taking advantage of Magaletta et al. dataset that covers late stages of pharyngeal development including early cTEC and mTEC even though no bona fide transitory TEP could be identified. Since cTECs are the natural and most direct product of TEP differentiation, we focused on cTEC as the target population for this step of differentiation. Hierarchical clustering identified a cluster of DOE samples that have the highest expression for markers of cTEC and also of its close mTEC population and of thyroid cells (**Figure 1D, Bottom Right**). Although our statistical approach could not discriminate cTEC from mTEC or thyroid for their resemblance to TEP, we took this population cluster as a distant proxy of TEP. The ANOVA on the mean expression of cTEC markers identified a statistically significant effect for the addition of CHIR99, as well as the deprivation of FLI06 and IWR1. Similarly to above, we chose to keep FGF10 and EGF since.

We summarized the results of the DOE-based screen with the pathways that needs to be activated (**+**), inhibited (**-**) or that shouldn’t be inhibited (**no sign**) to drive iPSc differentiation into TEPs (**Figure 1E)**. The pathways that we chose to activate by factors that we selected even though neutral in the screen are shown in blue.

### Characterization of the differentiation product demonstrates efficient differentiation into TEP

To follow the thymic differentiation process at the transcriptomic level, we sequenced samples at multiple time points across our differentiation protocol by bulk RNA-seq. Human primary TECs, sorted on CD45^-^EPCAM^+^ and originating from thymic samples of pediatric patients undergoing cardiac surgery, were included to assess the state of maturation of the differentiation product. iPSc samples were included as well as the starting point of the differentiation trajectory. We performed differential gene expression of each sample against iPSc and identified 9 gene modules enriched at distinct differentiation stages and comprising known markers of thymic differentiation stages such as *SOX17* (DE), *HOXA3* (AFE), *EYA1* (3PPE) or *EPCAM* (TEP) that hits in the same module than CXCL12 whose expression has been shown to be a common feature of cTECs and to attract DP thymocytes by for positive selection (Lucas et al., 2017) (**Figure 2A, Left**). However, markers associated with a more mature TEC phenotype, such as cortical proteasome components encoded by *PSMB11* and *PRSS16*, or genes involved in antigen presentation (*HLA-DRA*, *CD80*, *CD86*), were not expressed in late differentiation cultures and did not reach the expression level measured in primary TECs. To determine the biological processes enriched in the 9 gene modules, Gene Ontology (GO) terms were associated using ClusterProfiler. Ordering GO terms by time of differentiation showed a shift from pluripotency state and cell amplification to embryogenesis and organ morphogenesis (**Figure 2A, Right**). As expected, GO terms in late differentiation samples and primary TEC controls were associated with lymphocyte interaction and regulation of immune response, indicating the enrichment of our differentiation product in TEC-associated genes. Detailed list of GO terms is available in **Supp Figure 4**. Together, these results show the progressive loss of pluripotency and acquisition of TEC-fate through modulation of the pathways we identified by the DOE-based screen.

**Figure 4.**
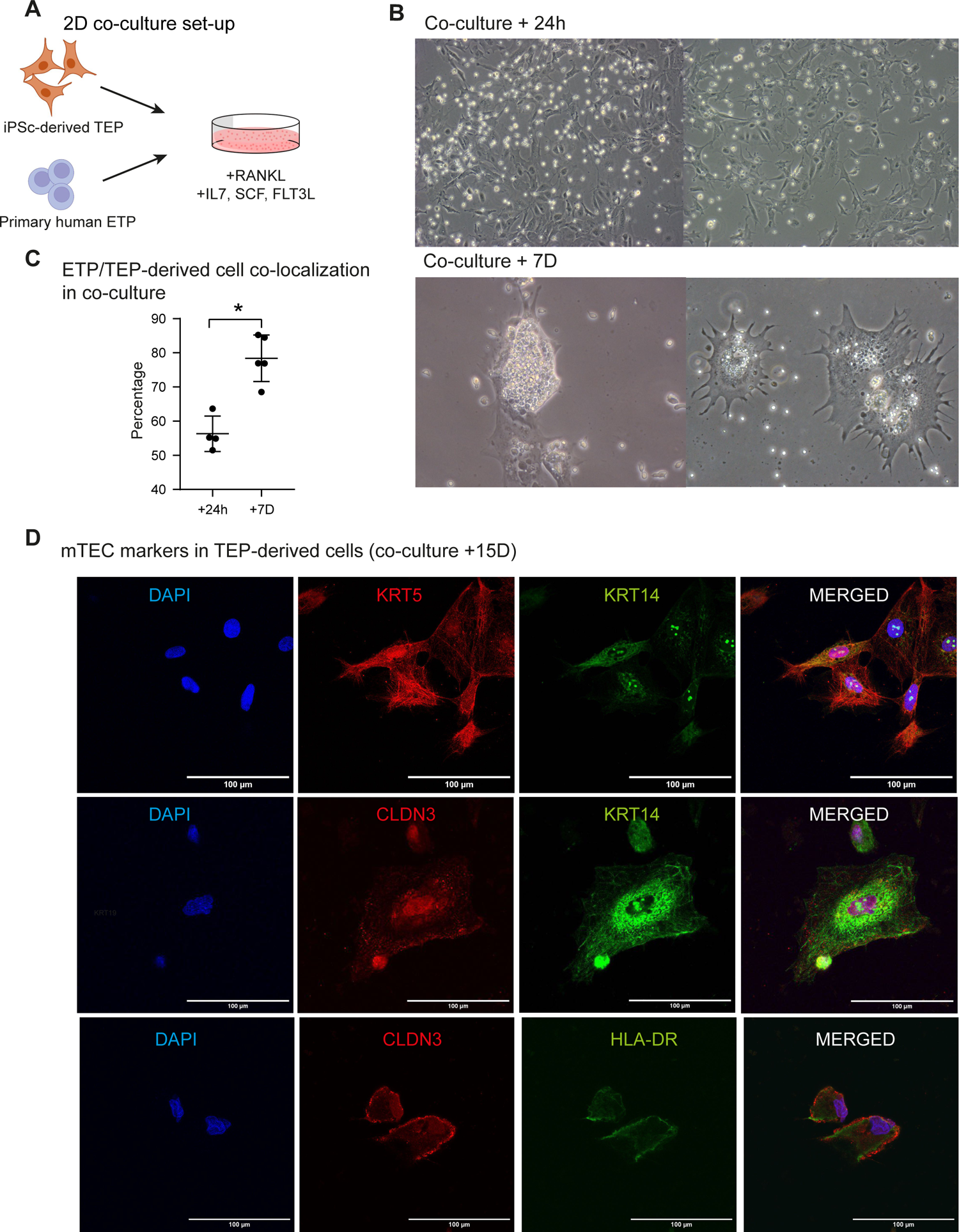
Early thymocytes (ETP) colocalize with TEPs in co-cultures. **(A)** 2D co-culture of D13 TEP differentiated from iPSc with primary human ETPs with RANKL, SCF, IL7 and FTL3L. **(B)** Bright field optical microscopy images of TEP coculture with ETP after 24h and 1 week of co-culture. **(C)** ETP/TEP co-localization quantification. **(D)** lmmunofluorescence by confocal microscopy of D17 TEPs co-cultured with ETPs of mature and medullary TEC markers. Nuclei are stained with DAPI.

Another post-DOE optimization was performed considering recent publications on thymic differentiation highlighting the benefits of FGF8 supplementation for AFE induction (Provin and Giraud, 2022). We confirmed similar results in our protocol by measuring a significant increase of *FOXN1* expression at the end of differentiation in samples exposed to 50 ng.mL^-1^ FGF8 from day 4 to 6 (**Supp Figure 5**).

**Figure 5.**
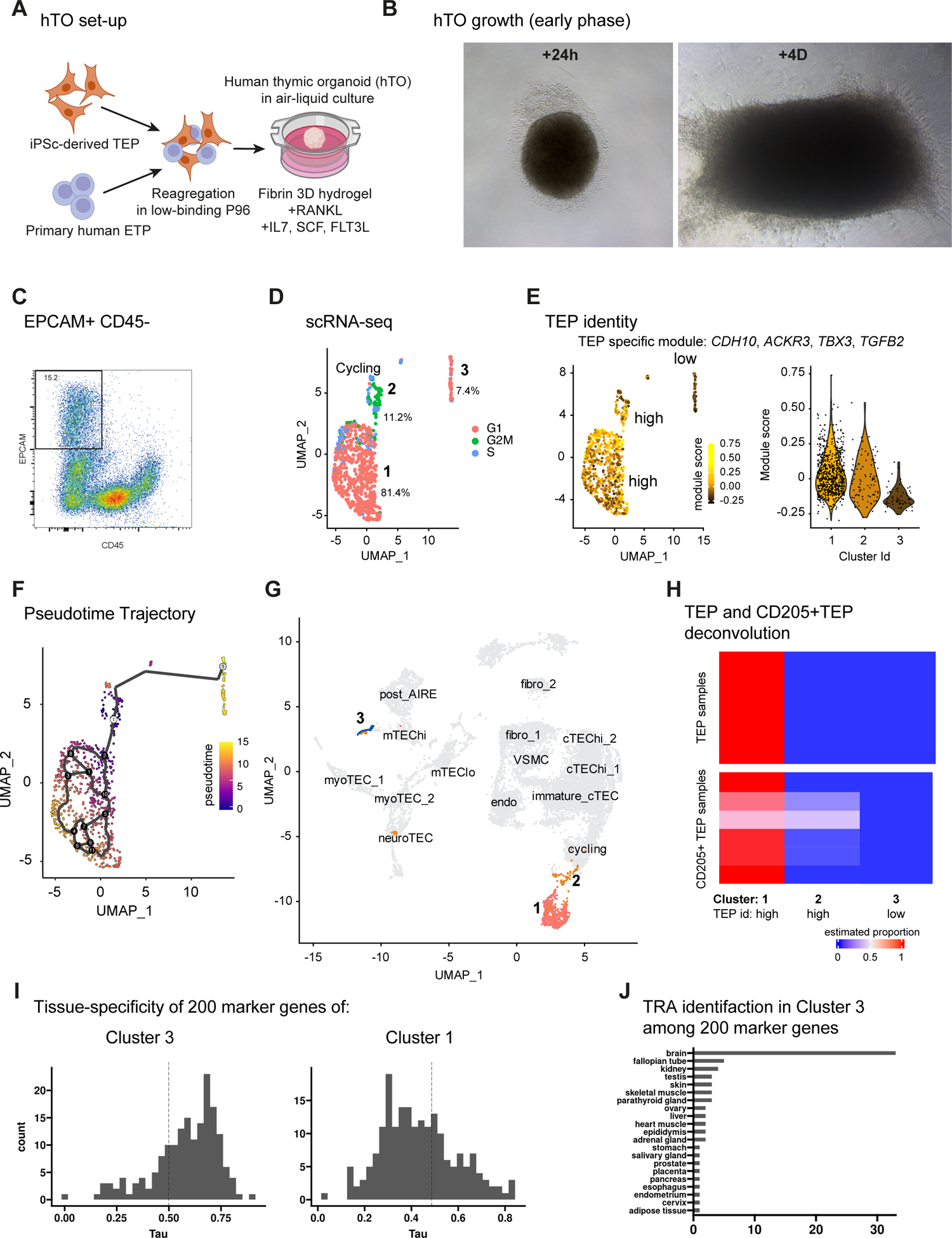
Early phase of hTO differentiation. **(A)** Reaggregation of D13 TEP differentiated from iPSc with primary human ETPs forms human Thymic Organoids (hTOs) cultured in a 3D fibrin hydrogel in air-liquid interface supplemented with RANKL, IL7, SCF and FTL3L. **(B)** Bright field images of hTO at 24h and 4 days. **(C)** Flow cytometry analysis of D7 organoids, gating on TEC (EPCAM^hi^CD45^-^). **(D)** scRNA-seq of EPCAM^hi^CD45^-^ cells and cell cycle scoring. **(E)** Module score of specific genes to iPSc-derived TEPs. **(F)** Pseudotime trajectory of EPCAM^hi^CD45^-^ cells with cluster 2 selected as root. **(G)** Integration of scRNA-seq data of EPCAM^hi^CD45^-^ cells (Clusters 1, 2 and 3) with a human thymic atlas. Gray is for cells from the atlas. **(H)** Deconvolution of TEP and CD205+TEP onto scRNA-seq of EPCAM^hi^CD45^-^ cells. **(I)** Distribution of the tissue-specific variable, Tau, within clusters 3 and 1. **(J)** Evaluation of the TRA content expressed in Cluster 3 with a surfeit of brain antigens.

Altogether, our findings led to put forward an optimized protocol of human iPSc differentiation into TEPs (**Figure 2B**). To finally assess its efficiency, we performed a deconvolution analysis of bulk RNA-seq data of the end-product using Magaletta dataset. Remarkably, this approach identified cTEC as the closest population from the reference to our differentiation product confirming we generated TEP with high efficiency (**Figure 2C**). In addition, we showed that all the generated cells stained positive for the thymic identity markers, PAX9 and FOXN1, revealing a complete differentiation towards TEC-fate (**Supp Figure 5**). We then benchmarked our protocol against two state-of-the-art protocols from Sun *et al*. and Parent *et al*. studies, by measuring the expression of *PAX9* and *FOXN1*. We found that our new protocol yielded cells expressing significantly higher levels of *PAX9* et *FOXN1* (**Supp Figure 5**). To test if those results came from a protocol overfitting to our iPSc cell line LON71, or a true optimization of the biological phenomenon, we differentiated two other iPSc cell lines (MIPS203 and LON80) and showed no significant differences in *FOXN1* and *PAX9* expression, excepted for the MIPS203 that showed lower *FOXN1* expression (**Supp Figure 5, E**).

### CD205+ TEPs are the main product of differentiation and are equipped for thymic crosstalk

Since it is well established that CD205+ TEPs are bipotent progenitors of mature cTECs and mTECs, we asked whether the iPSc-derived TEPs we generated are positive for the CD205 surface marker and in which proportion. We confirmed by flow cytometry that almost all cells expressed the epithelial marker EPCAM and found that a large majority of them were positive for CD205, as shown in **Figure 3A, Left**. Controls were also shown to attest the specificity of the staining (**Figure 3A, Middle**). To generalize this result, we performed a series of independent differentiation and assessed the proportions of EPCAM+ and CD205+ cells at the TEP but also at the earlier 3PPE stage for comparison. We found that the median percentage of cells that are positive for EPCAM at the TEP stage is close to 100% and those expressing the additional CD205 to 70% (**Figure 3A, Right**). As expected, CD205 expression was not detected at the 3PPE stage in contrast to EPCAM that marks the epithelial cell fate.

We then asked whether the bona fide CD205+ TEPs are equipped to sense maturation signals from the developing thymocytes and provide signals to the latter to promote their differentiation. To this end, we performed RNA-seq of the sorted CD205+ TEP populations and analyzed the expression of a set of genes encoding proteins involved in responsiveness of maturing TEC to signals provided by the developing thymocytes and in their ability to promote thymocyte development. We found a set of genes showing preferential expression in CD205+TEP and coding for proteins involved in the classical and alternative NF-κB signaling pathway with RELA and RELB constituting the NF-κB molecular complexes, TRAF5 mediating signal transduction, RANK (*TNFRSF11A*) receiving RANKL signals from thymocytes and activating the NF-κB pathway (**Figure 3B, Left**). Activation of the alternative NF-κB pathway is a key element for acquisition of a mTEC fate and features enabling mature mTEC to sustain self-antigen expression and presentation to developing thymocytes by MHCII molecules. We also identified the NOTCH ligands DLL1, DLL4, JAG1 and JAG2 whose coding genes show robust higher expression in TEP vs iPSc (**Figure 3B, Right**). Activation of the NOTCH pathway is crucial to promote thymocyte development. Interestingly, we observed that all or CD205+ TEPs have an enhanced expression of *KRT19* that has recently been shown to be a marker of medullary fate in CD205+ TEPs (Lucas et al., 2023). Together our findings indicate that the CD205+ TEPs that we generated are likely to sense signals susceptible to activate medullary differentiation programs in TEP.

### Expression of mature and medullary TEC markers in TEP-derived cells after 2D co-culture with ETPs

While the maturation into functional TEC *in vitro* has proved to be challenging, with most studies grafting TEP in mice (Gras-Peña et al., 2022; Parent et al., 2013; Ramos et al., 2022; Sun et al., 2013), we aimed at inducing TEP maturation fully *in vitro*. Thymocytes have been identified as a crucial source of signalization that regulates thymic medulla formation and maturation, especially for the mTEC^hi^ compartment (Lopes et al., 2015).

Therefore, we investigated the capacity of early thymocytes to promote TEP maturation. Early thymic progenitors (ETP) from human thymus samples were harvested and sorted by flow cytometry using CD3^-^CD4^-^CD8^-^CD14^-^CD19^-^CD56^-^(Lin)CD34^+^CD7^+^ (**Supp Figure 6**). This rare population constitutes multipotent hematopoietic progenitors upstream of the T cell differentiation (Deng et al., 2021; Lavaert et al., 2020). We characterized the sorted cell population by flow cytometry and validated their ETP phenotype (**Supp Figure 6**). Sorted ETP were CD45^+^CD3^-^CD34^+^, mostly double negative (DN) for CD4 and CD8, and expressed both CD62L and CD44. A minor CD44^-^ population could include already committed DN3 thymocytes.

**Figure 6.**
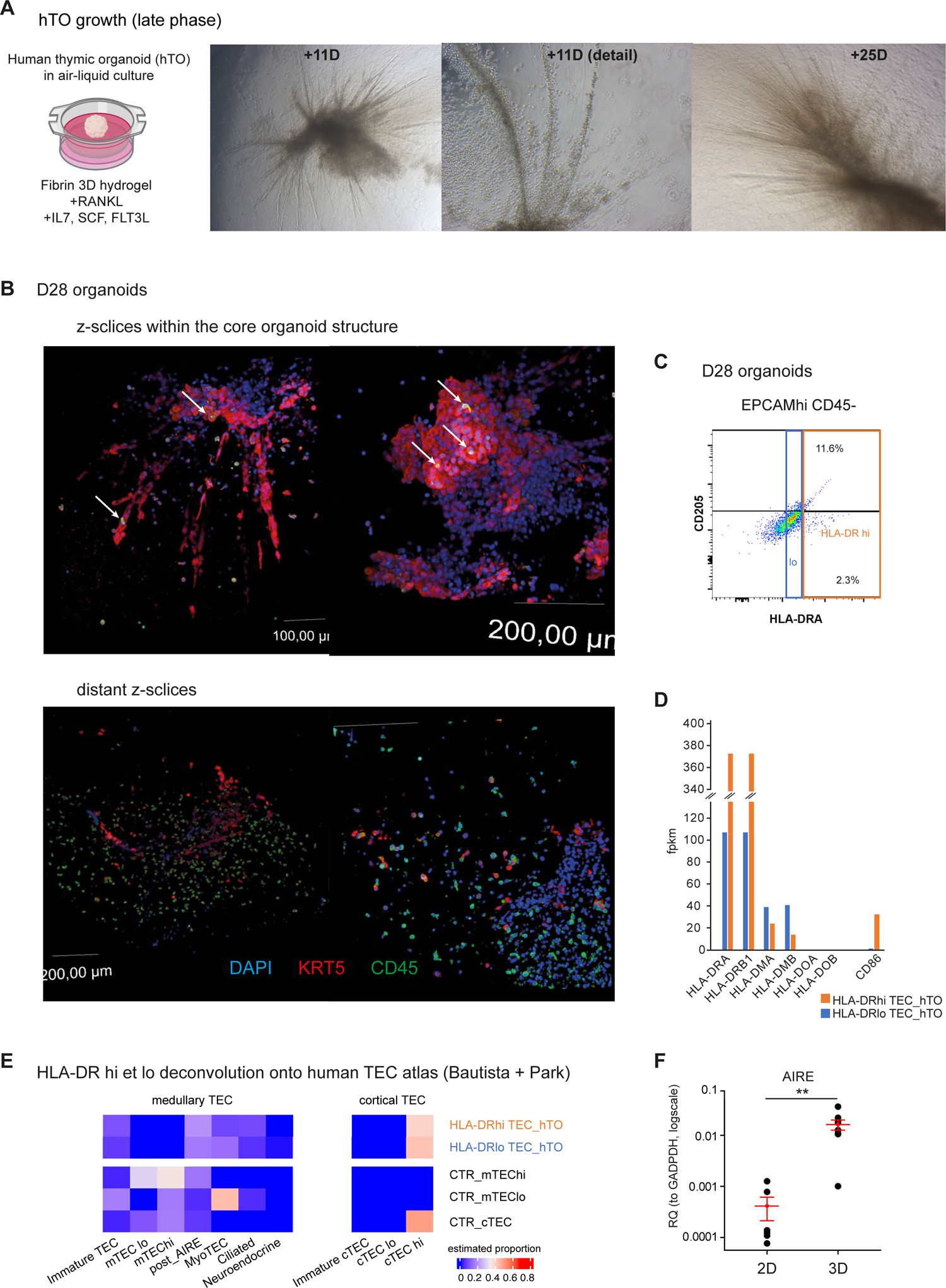
Late phase of hTO differentiation. **(A)** Bright field images of hTO at 11 and 25 days. **(B)** Flow cytometry analysis of D11 organoids, gating on TEC (EPCAM^hi^CD45^-^). **(C)** 3D immunofluorescence confocal imaging of the mature epithelial compartment (KRT5) and hematopoietic compartment (CD45) in D28 hTOs. **(D)** fpkm expression of genes encoding HLA-DR, DM, DO and the co-stimulatory CD86 molecule from bulk RNA-seq data. **(E)** Deconvolution of HLA-DRhi and HLA-DRlo TECs from hTOs onto a human thymic atlas. Control mTEChi, mTEClo and cTEC samples were isolated from human thymic samples and included. **(F)** qPCR quantification of AIRE in D23 TEP-ETP co-culture in hTO system (3D) and in classical monolayer culture (2D). Relative quantification to GAPDH, (n=6).

To test whether the TEPs we generated have potential to differentiate towards a medullary cell fate in addition to a cortical one, we perform simple 2D co-cultures of TEPs and primary ETPs in supplementing the medium with cytokines stimulating TEC, and T cell growth and maturation, namely, RANKL, IL7, SCF and FTL3L (Lavaert et al., 2020; Pinto et al., 2013). Interestingly, from a seemingly random repartition shortly after co-culture set up, ETPs nearly exclusively located at the surface of the large, differentiated epithelial cells after one week (**Figure 4B**). Quantification of the co-localized cells confirmed its significant increase after one week of co-culture (**Figure 4C**). Given the capacity of TECs to interact with thymocytes and even to incorporate them, forming thymic nurse cells (TNC) complexes, it is possible that this physical interaction is concurrent with cell-cell signalization during thymopoiesis. However, we did not investigate further if thymocytes were incorporated by TECs or only located at their surface.

Finally, we performed Immunofluorescence experiments and found surface expression of KRT5 and KRT14 (**Figure 4D**). In humans, *KRT14* is a specific marker of mTECs, whereas *KRT5* is expressed in all mature TECs with a preference for mTECs. The medullary feature of KRT14 cells was highlighted by positive co-staining with CLDN3, a marker of the most differentiated mTECs, detected with typical focal point staining. In addition, we found the presence of few cells positive for HLA-DR expression, some of them showing co-expression with CLDN3. Together these results show that the generated TEPs have the potential to differentiate into mature TECs and, for a fraction of them, to TECs positive for medullary markers, therefore strengthening the potential of the TEP we generated to be responsive to signals that drive medullary differentiation.

### An 3D thymic organoid system further promotes medullary differentiation

Classical culture in monolayer is known to be unsuitable for TEC maintenance, leading to a quick loss of TEC functional markers (Pinto et al., 2013). Considerable progress has been made recently, with development of 3D culture systems in hydrogels and the set-up of artificial thymic organoids (Campinoti et al., 2020; Fan et al., 2015; Hun et al., 2017; Montel-Hagen et al., 2020). We developed a human thymic organoid (hTO) coculture system by reaggregating human primary ETPs with the iPSc-derived TEPs and seeding the cell mass in a fibrin hydrogel (**Figure 5A**). Organoids were cultured in air-liquid interfaces on cell inserts, in the same XVIVO10 base culture medium, as other tested media such as DMEM did not supported comparable cell growth (data not shown). As for the 2D co-culture, we supplemented this medium with RANKL, IL7, SCF and FTL3L. After 24 hours, organoids showed a spheroid shape with a crown-like structure of early thymocytes (**Figure 5B Left**). Organoids showed signs of growth from day 4 after seeding, with total size increase and formation of cellular projections in the hydrogel (**Figure 5 Right**).

We took advantage of the fact that the cells composing the hTOs can be easily harvested with low mortality in the early phase of the hTO growth, to perform scRNA-seq experiments and monitor early TEP differentiation at day 7. We investigated by flow cytometry the cellular composition of the organoids and identified an EPCAM+CD45-population that likely corresponds to TEP-derived cells (**Figure 5C**). scRNA-seq of this population revealed one big cluster 1, a smaller cluster 2 close to the former and composed of cells showing cycling features, and a distant smaller cluster 3 (**Figure 5D**). To follow the fate of the TEP population in the organoids, we identified a set of 4 genes, *i.e.*, *CDH10*, *ACKR3*, *TBX3* and *TGFB2*, that are specifically expressed in iPSc-derived TEPs in comparison to primary TEC, to thymic precursor cell populations of the Magaletta dataset and to TEC mature sub-populations of the human thymus corresponding to the reference scRNA-seq dataset from the Human Protein Atlas (https://www.proteinatlas.org/). We computed the score of expression of the gene module and found strong expression in the main cluster 1 and in the cycling cells, revealing the strong TEP identity of these cells. In contrast, the more distant cluster 3 and a fraction of cycling cells showed low TEP identity, indicating that these cells may stem from the iPSc-derived TEPs in the organoids (**Figure 5E**). To test this hypothesis, we performed a trajectory analysis using Monocle3 by setting the cycling cells as the root of the trajectories. Remarkably, we found that the graph is fully connected showing that all cells are in the same partition and thus developmentally linked (**Figure 5F**). We identified a trajectory from the cycling cells to the cells with high TEP identity and a second, from the cycling cells to the low TEP identity cells.

To identify the nature of the low TEP identity cluster, we integrated the organoid scRNA-seq data to the human TEC atlas from Bautista et al. (Bautista et al., 2021). Remarkably, we found that the low TEP identity cells from cluster 3 were located within the medullary part of the UMAP, in close vicinity of mTEChi, whereas the high TEP identity cells from cluster 1 lied in the cortical part of the graph next to immature cycling cTEC (**Figure 5G**). In addition, the absence of detected expression of *HLA-DRA* or *KRT5* indicated that these cells still are at an immature stage.

To ensure that the low TEP identity cells arose after the organoid formation and were not already present in the iPSc-derived TEPs, we performed a deconvolution analysis of the bulk RNA-seq data of the iPSc-derived TEPs and the purified CD205+ TEPs on the three clusters identified in the organoid scRNA-seq. Remarkably, we found that the main high TEP identity cluster 1 was the closest to the iPSc-derived TEPs that were negative for a mixture with low TEP identify cells (**Figure 5H**). Comfortingly, we also showed that the sorted CD205+TEPs correspond to the high TEP identity clusters. Finally, we assessed the enrichment of TRA gene expression among the genes that are most specific to the low TEP identity cluster 3. To this end, we computed for each gene, the tissue specificity index, Tau, by implementing its calculation algorithm in a shiny program that we developed to parse the comprehensive human tissue-specific gene expression database, GTEx. Remarkably, we found a shift in the Tau distribution towards higher values in the low TEP identity cluster 3 in comparison to the cluster 1, with almost 40% of the 200 tested genes considered as tissue specific (Tau > 0.7). In addition, most of the identified TRA genes were of brain origin, consistent with the previous observation of a preferential expression of brain-specific TRAs in immature mTEC populations.

Together our findings show that upon 3D interaction with primary ETPs, the iPSc-derived TEPs start to differentiate towards a medullary fate in the early phase of the organoid growth and show feature of TRA-favored expression.

### TEC maturation in the late phase of orgnanoid differentiation

hTO could be maintained several weeks in the 3D culture system and reached 5 mm size. At day 11 organoids morphology evolves from a spheroid to a complex structure with multiple projections colonizing the gel (**Figure 6A**). Interestingly, thymocytes-like cells were concentrated around the cellular projections. To assess the level of TEC maturity using KRT5 and the interaction of these cells with thymocytes (CD45+) in the late phase of the organoid development, we performed confocal microscopy at day 28 (**Figure 6B, Top**). Remarkably, we observed extensive KRT5 staining of the cellular projections indicative of mature epithelial features in the organoids. Moreover, physical interaction between thymocytes and TECs in the organoids could be observed with the co-localization of CD45^+^ cells at the surface of the epithelial structures. Interestingly, distant z-slices from the organoid, in the hydrogel volume, showed dispersed thymocytes, highlighting their ability to freely circulate in the gel which was originally devoid of cells during its casting (**Figure 6B, Bottom Left**). This property was also observed for KRT5+ cells, with migration far from the initial seeded micromass (**Figure 6B**, **Bottom Right**).

We then investigated the cellular composition of D28 organoids by flow cytometry, and identified four main populations that likely correspond to the TEC compartment (EPCAM^+^CD45^-^), thymocytes (EPCAM^-^CD45^+^) and two uncharacterized EPCAM^-^CD45^-^ and EPCAM^lo^CD45^+^ populations (**Figure 6C**). Further analysis of the EPCAM^+^CD45^-^ compartment composition, relative to HLA-DR and CD205 expression, revealed a global decrease of CD205 and a set of cells that showed strong expression of HLA-DR. To characterize this population further, we isolated the HLA-DR-high and low cells and perform highly sensitive bulk RNA-seq. We found strong expression of HLA-DR encoding genes, namely, *HLA-DRA* and *HLA-DRB1*, with higher levels in HLA-DR high than low cells, confirming expression of the HLA-DR complex in the organoid at day 28 (**Figure 6D**). Remarkably, HLA-DR high cells showed a solid expression of *CD86* that encodes an important costimulatory molecule for the priming and activation of naive and memory T cells. In addition, we found that the genes coding for HLA-DM but not HLA-DO were also expressed indicating that the HLA-DR high cells are able to present processed self-antigen peptides through MHCII molecules, and provide activation signals to developing thymocytes, therefore recapitulating antigen presenting cell features of mature TECs.

Finally, to evaluate the cellular composition of the HLA-DR high and low cell fractions we performed deconvolution onto the human TEC atlas as an alternative to scRNA-seq and direct cell population identification. Indeed, harvesting the tightly imbricated cells from late organoids suffers poor recovery and high mortality, therefore hindering scRNA-seq approaches. Remarkably, inference of the relative abundance of the known human TEC sub-populations in the HLA-DR high and low TEC samples revealed a mixture of cells that include terminally differentiated mTEC populations including corneo-like mTECs, myoid cells and ciliated cells also referred as mTEC mimetic cells, as well as cTEC mature populations (**Figure 6E**). The absence of immature TEC populations indicates that the iPSc-derived TEP we used to form the organoids kept differentiating after day 7 and lost their immature phenotype. However, despite the observation of higher levels of *AIRE* expression by qPCR in comparison to the 2D co-culture system (**Figure 6F**), we don’t detect its transcripts in RNA-seq data nor hints of bona fide mTEChi by deconvolution, suggesting that *AIRE* expression might not reach levels compatible with its transactivation function or desynchronized in the maturing TECs.

Together, these findings showed differentiation/maturation in the late phase of organoid growth with TEC acquiring mature medullary and cortical features, as well as the capacity to present self-antigen peptides to developing thymocytes through high MHCII expression.

### hTOs support thymopoeisis and maturate into SP CD4+ and CD8+ T lymphocytes with thymic emigrant phenotypes

To investigate the capacity of hTOs to differentiate ETPs into mature SP T cells, hTOs were dissociated and analyzed by flow cytometry. Soft mechanical dissociation yielded mostly non-adherent CD45+ hematopoietic cells. Expression of CD34 was absent, confirming the loss of the hematopoietic progenitor identity, and thus ETP differentiation (**Supp Figure 7**). Importantly, we observed a CD3+ population arising at D17 and rapidly enlarging at W3 to reach up to 60% at W5, therefore confirming the commitment to a T cell fate (**Figure 7A, B**). TCR expression was analyzed by flow cytometry to decipher αβ vs γδ T lineage orientation in hTO (**Figure 7C**). Both αβ and γδ populations were detected in CD3^+^ thymocytes. αβ lineage was nonetheless predominant in comparison to rare γδ thymocytes, in contrast to recent results using a similar system (Hosaka et al., 2021).

**Figure 7.**
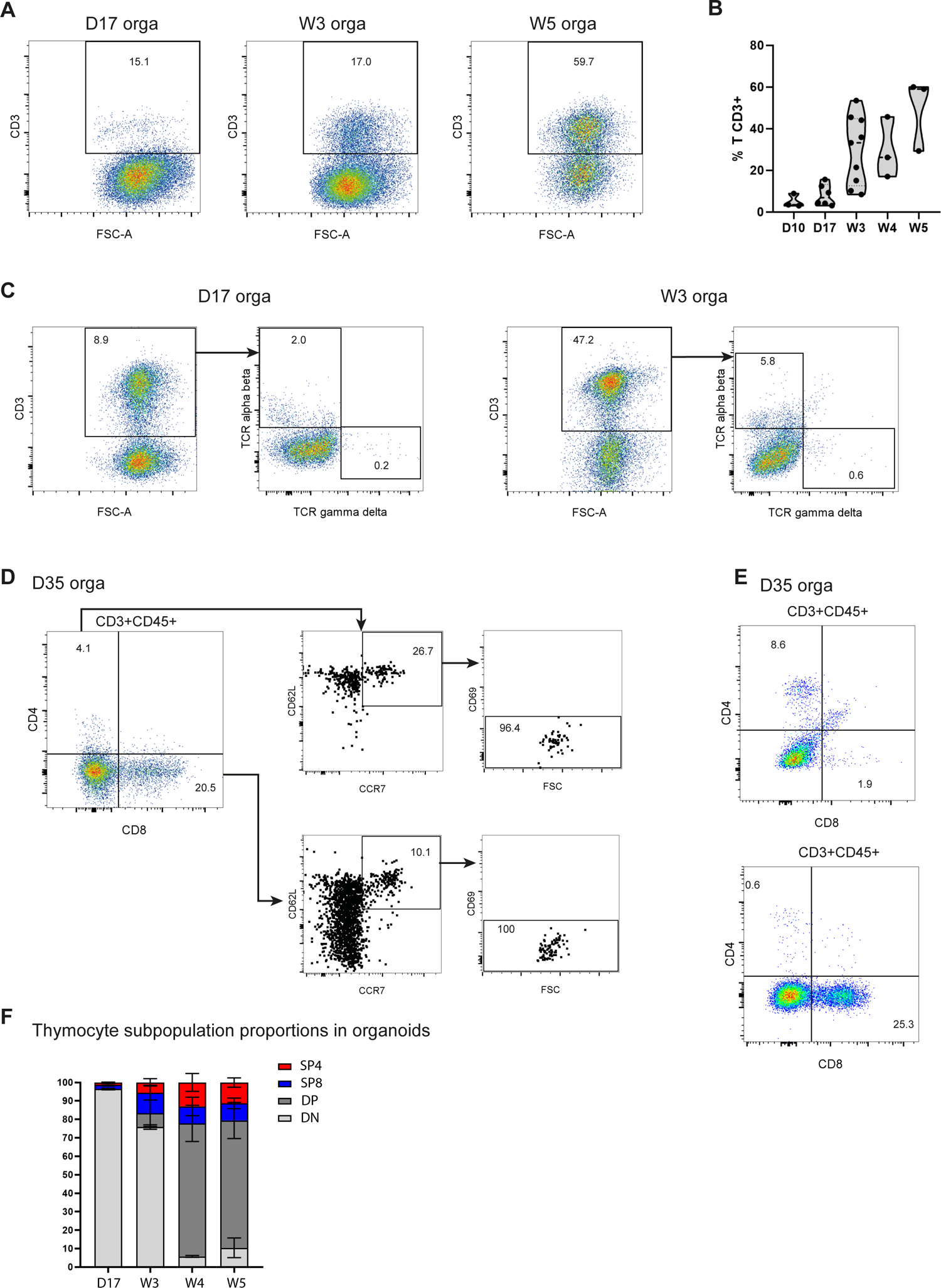
hTO induce ETP maturation into SP CD4+ and CD8+ T lymphocytes with thymic emigrant phenotypes. **(A, B)** Flow cytometry analysis of the kinetic of thymopoeisis in hTOs shows accumulation of CD3+ T cells, with extensive increase after week 3 of co-culture. **(C)** Flow cytometry expression of both alpha beta and gamma delta TCRs in W3 hTOs. **(D)** Flow cytometry of the hematopoietic fraction of D35 hTO shows presence of SP CD8+ T cells and less abundant SP CD4+ T cells. A subset of SP populations shows a mature phenotype CD62L^+^CCR7^+^CD69^-^ typical of thymic emigrants. **(E)** Variability in the relative frequency of generated SP CD8+ and SP CD4+ T cells. **(F)** Summarization of thymocyte population proportions obtained in a number of independent hTO experiments, (n=6).

Remarkably, although most of the CD3^+^ cells are CD4^-^CD8^-^ DN thymocytes, we found that hTOs also include CD4^+^CD8^+^ DP, CD4^+^CD8^-^ SP4 and CD4^-^CD8^+^ SP8, albeit in lesser proportions (**Figure 7D**). In this hTO experiment, a prominent SP CD8+ population along with a smaller SP CD4+ population were detected, revealing SP CD8+ and CD4+ thymocyte differentiation and showing a preferential maturation towards the CD8 fate. However, we observed experimental variability of the generated SP CD4^+^ and CD8^+^ T cell frequencies, with some hTOs showing higher proportions for CD4+ (**Figure 7E**). SP CD4+ thymocyte generation reflects a MHCII-dependent selection inside hTOs, and SP CD8+/CD4+ heterogeneity, a potential variation of MHCII expression by TEC populations. Importantly, generalization on multiple independent hTO experiments showed an equivalent ∼10% proportion of the generated SP CD8+ and SP CD4+ T cells (**Figure 7F**). We then assessed the state of maturation of the SP cells by quantitating the expression of the markers CCR7, CD62L and CD69. Remarkably this analysis revealed minor populations of SP T cells in the hTOs that displayed mature CCR7^+^CD62L^+^CD69^-^ phenotypes, demonstrating the hTO capacity to drive thymopoiesis, especially considering that ETPs cultivated alone showed high mortality (**Supp Figure 7**).

### scRNA-seq demonstrates multilineage differentiation of dendritic and mature T cells

To characterize in more details the CD45+EPCAMlo compartment of hTOs, we performed highly sensitive bulk RNA-seq on the sorted population (**Figure 8A**). Gene ontology analysis of differentially expressed genes relative to iPSc showed enrichment of terms linked to antigen processing and presentation by the MHCII complex, as well as to interaction with lymphocytes (**Figure 8B**). In addition, deconvolution analysis of the CD45+EPCAMlo transcriptome onto the thymus single cell atlas from Park et al. (Park et al., 2020) which includes the CD45+ cell fraction, showed that dendritic cells 2 (DC2) most likely constitute the main part of the CD45+EPCAMlo population (**Figure 8C**). Noteworthy, one of the two tested CD45+EPCAMlo samples shows slight positivity for DC1, indicating that CD45+EPCAMlo could fit a mixture of DCs including an important proportion of DC2 and potentially a small number of DC1.

**Figure 8.**
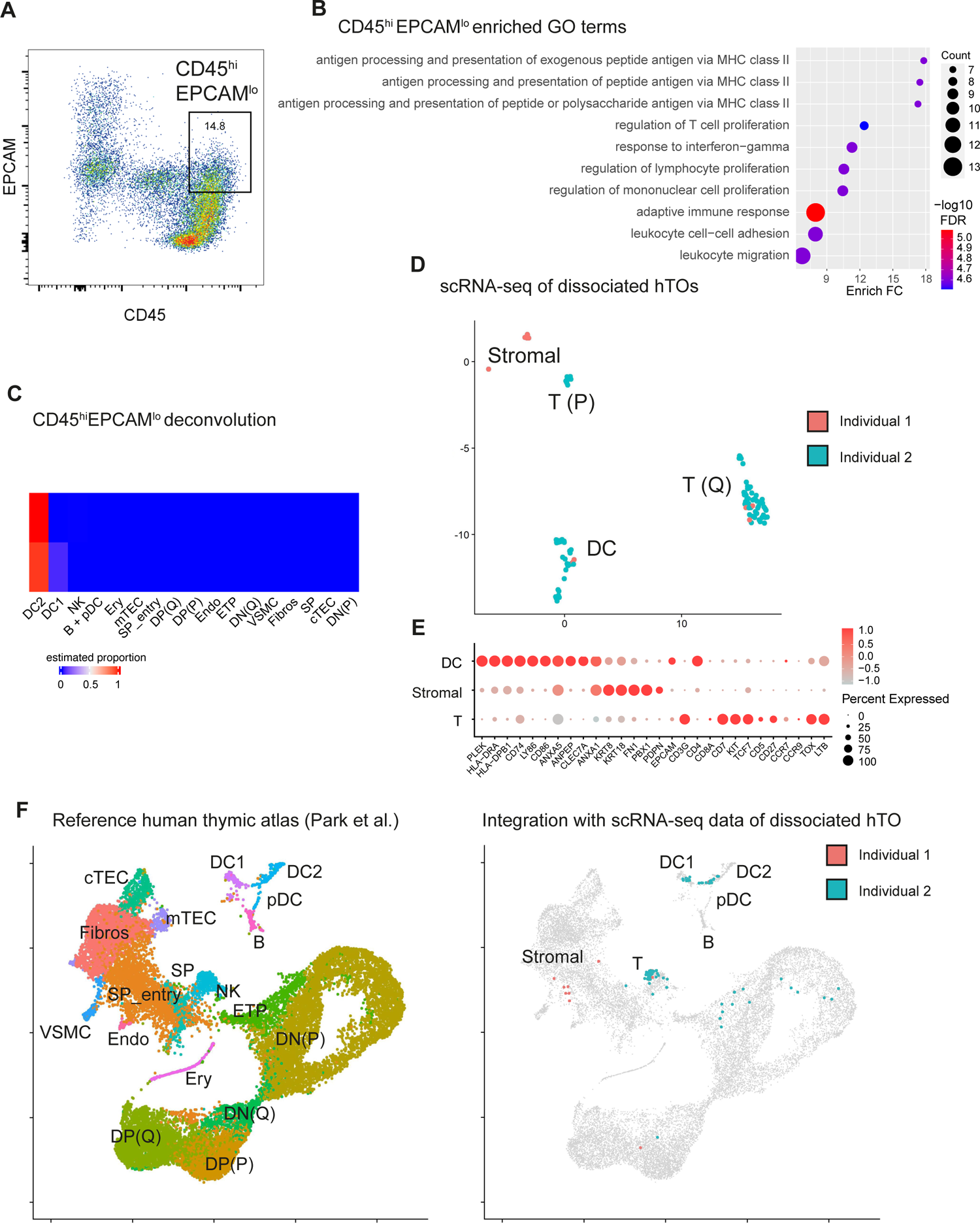
single cell RNA sequencing (scRNA-seq) of hTOs demonstrates multilineage differentiation of dendritic cells and mature T lymphocytes. **(A)** The CD45^hi^EPCAM^lo^ population in D28 hTOs was sorted by flow cytometry and their transcriptome sequenced by SMART-seq **(B)** GO terms in differentially expressed genes of CD45^hi^EPCAM^lo^ samples show enrichment of biological processes linked to antigen processing and presentation **(C)** Park et al. Thymic human cell atlas was reanalyzed and markers of the 3 main populations of thymic dendritic cells were identified. These markers are significantly upregulated in the CD45^hi^EPCAM^lo^ population transcriptomes, confirming their identification as dendritic cell. iPSc and differentiated TEPs were included as controls **(D)** D28 hTO hematopoietic compartment was further analyzed by scRNA-seq. UMAP projection reveals 3 main cell clusters. Cell origin, either from iPSc differentiation product or ETP maturation product, was assessed by comparing SNP profiles using scSPLIT, identifying the 2 individuals of origin and confirming that the DC and T population derive from a common progenitor. Cluster cell identity was determined by the expression of differentially expressed markers **(E)** CD3 and CD7 identify T lymphocytes, LY86 and PLEK the dendritic cell (DC) population and KRT8-18 a rare stomal population **(F)** To confirm cell cluster identification, data were projected on the thymic human atlas dataset from Park et al. using Seurat default integration on a reference data. **(G)** Our query data show prediction for SP T lymphocytes and DC clusters on the reference dataset.

To further explore the heterogeneity of this population and characterize the state of maturation of thymocytes in hTOs, we performed scRNA-seq of cells harvested from W4 by soft mechanical dissociation for CD45+ cell isolation. Three clusters were identified as illustrated by UMAP projection (**Figure 8D**). Two of them were of hematopoietic identity, as defined by *CD45* expression, whereas the third and minor one was negative for *CD45* consistent with a stromal identity. Differential gene expression pinpointed *KRT8*, *KRT18*, *FN1* and *PDPN* as markers of the stromal cluster, indicative of a TEP-derived origin. Identification of key markers for each hematopoietic cell cluster allowed identification of a dendritic (*PLEK*, *LY86*, *HLA-DRA*) and T cell population (*CD3E*, *CD7*, *TCF7*) (**Figure 8E**). In addition, T cells are divided in an actively proliferating minor cluster “T(P)”, and a main quiescent one “T(Q)” (**Figure 8D**). To finely label clusters and identify their *in vivo* counterparts, we projected our dataset as query to the reference thymic atlas from Park *et al*. (Park et al., 2020). Label transfer revealed high prediction scores for two main DC populations (DC1 and DC2) and SP mature T cells (**Figure 8F**) confirming that hTOs support multilineage hematopoietic differentiation. We note that the few stromal harvested cells showed weaker prediction scores indicating that these cells do not have any counterpart in the atlas reference. Indeed, besides the comprehensive capture of the CD45+ fraction, the reference only features two main homogeneous mTEC and cTEC clusters lacking the large diversity of TEC subpopulations. As expected, single cell transcriptomic data confirmed the maturation state of the generated T cells, with expression of markers such as *CCR7*, *CD27* or *IL2RA* (CD25) (**Figure 8E, Supp Figure 8**). To confirm that the DCs stemmed from ETPs and not from iPSc differentiation, we performed a single nucleotide polymorphism (SNP) analysis by using scSPLIT on the dissociated hTO scRNA-seq data. Each cell SNP profile was compared and regrouped by their individual of origin, the ETP donor or the donor from which iPSc were derived. As expected, the SNP analysis confirmed the cellular origin from two individuals, one corresponding to the stromal cluster and the other to the hematopoietic compartment (**Figure 8.D**). Therefore, both DC and T populations generated in the hTO originated from ETPs, revealing hTO ability to support multilineage hematopoietic differentiation.

## Discussion

This work provides a new protocol for *directed* differentiation of iPSc into TEP and their further maturation into functional thymic tissue supporting full thymopoiesis *in vitro*. Our original approach relies on the combination of an experimental design-based optimization of differentiation factor and the setup of a 3D organoid model leveraging thymocyte crosstalk in a three-dimensional hydrogel structure.

Previous proposed thymic differentiation protocols were built by testing only a few distinct combinations of factors, often iteratively, and without a robust statistical approach (Gras-Peña et al., 2022; Inami et al., 2011; Parent et al., 2013; Ramos et al., 2022; Soh et al., 2014; Sun et al., 2013). If this methodology can improve differentiation efficiency, it could nonetheless lead to suboptimal results or even confounding factor effects. A DOE-based optimization approach allows for less bias, by simultaneously testing multiple factor combinations. Moreover, readout quality is crucial for the optimization. Instead of relying only on expression of a few markers, we leveraged bulk RNA-seq and public scRNA-seq data from thymic and pharyngeal organogenesis to classify the efficiency of the differentiation. This multidimensional readout allowed us to measure the effect of each differentiation factor on our sample transcriptomes and to select those with highest similarity to *in vivo* developing pharyngeal populations. Although public data used as reference is from mice, thymus organogenesis was found to be mostly conserved with humans (Li et al., 2021). Moreover, the positive results of the optimization advocate for the pertinence of the approach. Because we used only one iPSc line for DOE optimization, there was a risk of overfitting the protocol to this specific line. However, we showed that our protocol was also able to differentiate two more iPSc cell lines into thymic tissue, demonstrating its robustness.

In addition to previously identified regulators of AFE and 3PPE induction, such as RA and WNT, we identified for the first time a positive effect of IGF1 and FGF10 supplementation on TEP differentiation. Conversely, our study contradicts previous reports in which BMP4 was used for AFE induction (Parent et al., 2013; Ramos et al., 2022). Indeed, we showed no significant effect of BMP4 nor the benefits of its inhibition by NOGGIN. Hedgehog inhibition showed no significant effect in our data as well. However, recent studies indicate that a fine temporal modulation of SHH promotes thymic development (Gras-Peña et al., 2022; Ramos et al., 2022), thus suggesting testing SHH activation or inhibition at other time points. Finally, our work suggested that systematic robust experimental design is an adapted tool for iPSc differentiation study, because of their inherent variability, temporal susceptibility, multifactorial aspect and required workload. Broader DOE designs, testing more factors with more informative readouts such as single-cell-OMICS could be powerful tools to decipher the gene regulation during cell differentiation. In addition, we reported high variability both inter- and intra-experiments, with variable differentiation yield. iPSc state during seeding could be an important source of variation. Reactive variability such as Matrigel is another potential source that must be carefully controlled. Thus, rigorous standardization and quality approach is required for reliable iPSc thymic differentiation.

Characterization of the differentiation product showed expression of TEP markers both at the RNA and protein level. However, weak expression of TEC markers demonstrates their immaturity, concordant with previous results where TEP grafting *in vivo* is required for the acquisition of functionality (Parent et al., 2013; Sun et al., 2013). However, development of an accessible and practical platform for thymopoiesis study and compatibility with future clinical applications imply an integral *in vitro* culture. A crucial contribution of this work is the setup of a 3D coculture system to mature TEP into HLA-DR^hi^ mTECs. Thymic crosstalk has been shown to be crucial for mTEC maturation and medulla structuration through signaling provided by thymocytes and involving RANK-LTB-CD40 pathways. Then, classical 2D culture has been shown to be detrimental to primary TEC functionality, while 3D culture promoted maintenance and maturation of primary TECs (Pinto et al., 2013). Therefore, we adapted previous works (Montel-Hagen et al., 2020; Pinto et al., 2013) to develop a fibrin-based hydrogel, seeded by ETP-TEP reaggregates. This resulted in the formation of hTOs growing at the air-liquid interface in 3D hydrogels. Since *in vivo* thymopoiesis relies on the successive circulation through cortical and medullary niches, a proper cortico-medullary segregation in hTOs must be pursued. Modulating matrix physical properties or avoiding the damaging TEP reaggregation stage could be promising approaches to reach this goal.

Although we did not report direct proof of thymic selection occurring in hTOs, we observed generation from ETPs of SP T lymphocytes with mature phenotypes CD62L^+^CCR7^+^CD69^-^. Because acquisition of those markers and survival signals is TEC-dependent on TECs, this demonstrates indirectly the hTO ability to perform thymic selection. Presence of SP CD4^+^ T lymphocytes in roughly similar proportion that SP CD8+ T cells demonstrate that selection of DP thymocytes into the CD4^+^ fate occurs in hTOs. Interestingly, TCR type analysis by flow cytometry showed both αβ and γδ T cells, confirming the ability of hTOs to support thymopoiesis, in contrast to studies showing abnormal exclusive γδ T generation (Hosaka et al., 2021). Interestingly, deeper analysis of the hTO hematopoietic compartment by scRNA-seq revealed the presence of dendritic cells (DC). This population can be linked to the EPCAM^lo^CD45^+^ population in hTO flow cytometry data. Given the recent reports revealing the crucial role of intrathymic DCs for thymocyte complementary selection, this unexpected population could participate in thymopoiesis in hTOs, for instance in promoting commitment to the SP CD4+ fate.

Our study thus provides insights on the regulation of iPSc thymic differentiation and advocates for the application of robust statistical tools to this subject. It provides an iPSc-derived human thymic organoid model, developed integrally *in vitro*. This multilineage organoid includes TEC, thymic fibroblasts, DC and T cells, and demonstrates thymic functionality, *i.e.,* the ability to generate mature SP T lymphocytes. Thus, it provides crucial resources for modeling thymopoiesis *in vitro.* Hence, use of iPSc opens promising long-term perspectives for regenerative medicine and cellular therapies with *in vitro* generation of engineered T lymphocytes.

## Acknowledgments

This work was supported by the EJP-Rare Disease JTC2019 program TARID project (ANR-19-RAR4-0011-05), la “Région Pays de la Loire “ (grant Bioregate) and a Novartis grant to MG. NP was supported by “la fondation d’entreprise ProGreffe”.

