## Supplementary Figures for "Combinatory differentiation of human induced pluripotent stem cells generates thymic epithelium that supports thymic crosstalk and directs dendritic- and CD4/CD8 T-cell full development"

Supp Figure 1

**A**

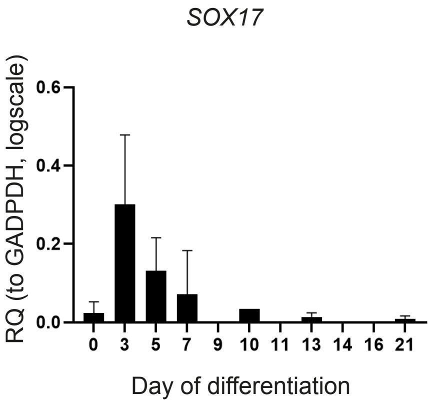


**B**

iPSc-derived cells (D5)

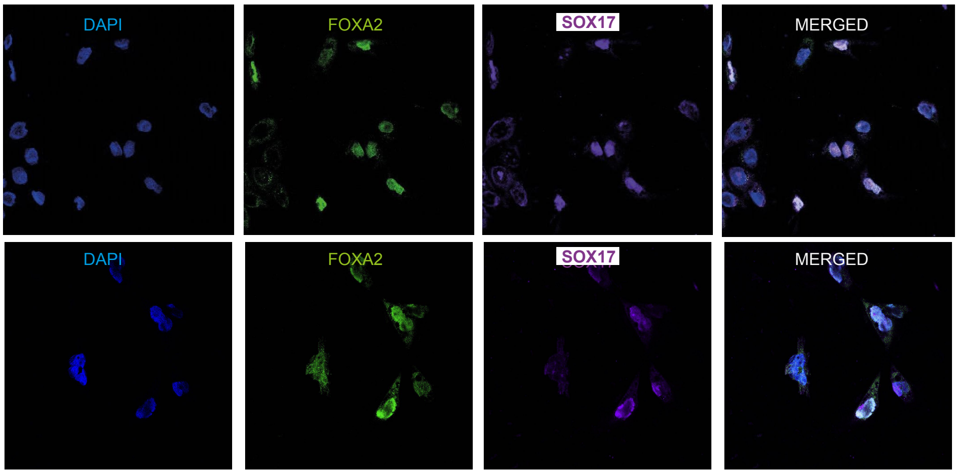


**C**

| Factor | Timing | Low dose | High dose | |
|---|---|---|---|---|
| RA | D5 - D7 | 0.075 | 0.75 | µM |
| LY3 | D5 - D7 | 0.1 | 10 | µM |
| IWR1 | D5 - D7 | 0.075 | 7.5 | µM |
| BMP4 | D5 - D7 | 0.01 | 10 | ng/mL |
| NOGGIN | D5 - D7 | 1 | 100 | ng/mL |
| BMP4 | D7-D11 | 0 | 10 | ng/mL |
| FGF8 | D7-D11 | 0 | 20 | ng/mL |
| IWR1 | D7-D11 | 0 | 7.5 | µM |
| FLI06 | D7-D11 | 0 | 5 | µM |
| CHIR99 | D7-D11 | 0 | 5 | µM |
| cyclo | D7-D11 | 0 | 1 | µM |
| BMP4 | D11-D13 | 0 | 50 | ng/mL |
| FGF10 | D11-D13 | 0 | 10 | ng/mL |
| IGF1 | D11-D13 | 0 | 10 | ng/mL |
| RANKL | D11-D13 | 0 | 50 | ng/mL |
| EGF | D11-D13 | 0 | 10 | ng/mL |

**D**

AFE: D5 - D7

| Run | RA | LY3 | IWR1 | BMP4 | NOG |
|---|---|---|---|---|---|
| 1 | 1 | 1 | 1 | -1 | -1 |
| 2 | -1 | -1 | 1 | -1 | -1 |
| 3 | 1 | -1 | -1 | 1 | -1 |
| 4 | 1 | -1 | 1 | -1 | 1 |
| 5 | -1 | 1 | 1 | -1 | -1 |
| 6 | -1 | -1 | 1 | 1 | 1 |
| 7 | 1 | 1 | -1 | -1 | 1 |
| 8 | -1 | 1 | -1 | 1 | 1 |
| 9 | -1 | 1 | -1 | -1 | 1 |
| 10 | 1 | 1 | 1 | 1 | 1 |
| 11 | -1 | 1 | -1 | 1 | -1 |
| 12 | -1 | -1 | -1 | -1 | -1 |
| 13 | 1 | -1 | -1 | -1 | -1 |
| 14 | 1 | 1 | -1 | 1 | 1 |
| 15 | 1 | -1 | -1 | 1 | -1 |
| 16 | -1 | -1 | -1 | 1 | -1 |
| 17 | -1 | -1 | 1 | -1 | 1 |
| 18 | -1 | -1 | 1 | -1 | 1 |
| 19 | 1 | 1 | -1 | -1 | 1 |
| 20 | -1 | -1 | 1 | -1 | -1 |
| 21 | 1 | -1 | 1 | 1 | 1 |
| 22 | 1 | -1 | 1 | 1 | 1 |
| 23 | -1 | -1 | -1 | -1 | 1 |
| 24 | 1 | -1 | -1 | -1 | 1 |
| 25 | -1 | 1 | 1 | 1 | 1 |
| 26 | 1 | -1 | -1 | 1 | -1 |
| 27 | -1 | 1 | 1 | 1 | -1 |
| 28 | 1 | 1 | 1 | 1 | -1 |
| 29 | 1 | 1 | -1 | -1 | -1 |
| 30 | -1 | -1 | -1 | 1 | 1 |
| 31 | 1 | 1 | 1 | -1 | 1 |
| 32 | 1 | 1 | 1 | -1 | -1 |
| 33 | -1 | 1 | -1 | -1 | -1 |
| 34 | 1 | -1 | 1 | 1 | -1 |
| 35 | -1 | 1 | 1 | 1 | -1 |
| 36 | -1 | 1 | -1 | 1 | 1 |

3PPE: D7 - D11

| Run | BMP4 | FGF8 | IWR1 | CHIR | FLI06 | cyclo |
|---|---|---|---|---|---|---|
| 1 | -1 | 1 | 1 | 1 | 1 | 1 |
| 2 | 1 | -1 | -1 | -1 | -1 | 1 |
| 3 | -1 | 1 | 1 | 1 | -1 | -1 |
| 4 | 1 | -1 | 1 | 1 | -1 | -1 |
| 5 | -1 | -1 | 1 | -1 | -1 | 1 |
| 6 | 1 | -1 | -1 | 1 | -1 | -1 |
| 7 | 1 | -1 | 1 | 1 | 1 | -1 |
| 8 | -1 | 1 | -1 | -1 | 1 | -1 |
| 9 | -1 | -1 | 1 | -1 | 1 | 1 |
| 10 | 1 | 1 | -1 | -1 | 1 | -1 |
| 11 | -1 | 1 | 1 | 1 | -1 | -1 |
| 12 | 1 | -1 | -1 | -1 | 1 | 1 |
| 13 | -1 | -1 | -1 | -1 | 1 | -1 |
| 14 | -1 | -1 | -1 | 1 | -1 | 1 |
| 15 | 1 | 1 | -1 | -1 | 1 | -1 |
| 16 | 1 | -1 | 1 | 1 | 1 | -1 |
| 17 | -1 | -1 | 1 | 1 | 1 | 1 |
| 18 | 1 | 1 | 1 | 1 | -1 | 1 |
| 19 | -1 | 1 | 1 | -1 | -1 | 1 |
| 20 | 1 | -1 | -1 | 1 | -1 | 1 |
| 21 | 1 | -1 | -1 | 1 | -1 | -1 |
| 22 | 1 | 1 | -1 | 1 | 1 | 1 |
| 23 | -1 | -1 | 1 | -1 | 1 | -1 |
| 24 | -1 | 1 | -1 | 1 | 1 | -1 |
| 25 | 1 | 1 | -1 | -1 | -1 | 1 |
| 26 | -1 | -1 | -1 | -1 | -1 | -1 |
| 27 | -1 | -1 | 1 | -1 | -1 | -1 |
| 28 | -1 | -1 | -1 | 1 | 1 | 1 |
| 29 | 1 | 1 | 1 | -1 | -1 | 1 |
| 30 | 1 | 1 | 1 | -1 | -1 | -1 |
| 31 | -1 | 1 | -1 | -1 | -1 | -1 |
| 32 | -1 | 1 | -1 | 1 | 1 | 1 |
| 33 | 1 | 1 | 1 | -1 | 1 | 1 |
| 34 | 1 | -1 | 1 | -1 | 1 | 1 |
| 35 | 1 | 1 | 1 | 1 | 1 | -1 |
| 36 | -1 | 1 | -1 | 1 | -1 | 1 |

TEP: D11 - D13

| Run | BMP4 | FGF10 | IGF1 | RANKL | EGF |
|---|---|---|---|---|---|
| 1 | 1 | 1 | -1 | -1 | 1 |
| 2 | 1 | 1 | 1 | -1 | 1 |
| 3 | -1 | -1 | -1 | -1 | 1 |
| 4 | 1 | 1 | -1 | -1 | 1 |
| 5 | 1 | -1 | -1 | 1 | 1 |
| 6 | -1 | 1 | -1 | 1 | -1 |
| 7 | 1 | 1 | 1 | 1 | -1 |
| 8 | 1 | -1 | -1 | -1 | -1 |
| 9 | -1 | -1 | -1 | -1 | -1 |
| 10 | -1 | -1 | -1 | 1 | 1 |
| 11 | -1 | 1 | 1 | 1 | 1 |
| 12 | -1 | 1 | -1 | -1 | -1 |
| 13 | -1 | 1 | 1 | -1 | -1 |
| 14 | -1 | -1 | 1 | -1 | 1 |
| 15 | 1 | 1 | -1 | 1 | -1 |
| 16 | 1 | -1 | 1 | -1 | 1 |
| 17 | 1 | -1 | -1 | 1 | -1 |
| 18 | 1 | -1 | 1 | -1 | -1 |
| 19 | -1 | 1 | 1 | -1 | -1 |
| 20 | -1 | 1 | -1 | 1 | 1 |
| 21 | 1 | -1 | 1 | 1 | -1 |
| 22 | 1 | 1 | 1 | 1 | 1 |
| 23 | -1 | -1 | 1 | 1 | -1 |
| 24 | -1 | -1 | 1 | 1 | 1 |

**Supplementary Figure 1.**
(A) State-of-the-art DE induction protocol results in transient peak of expression of SOX17 : qPCR relative quantification to GAPDH, (n=5).
(B) Proteic co-expression of DE markers FOXA2 and SOX17 at D5 by IF
(C) Factor abbreviations, doses and supplementation timings for the Experimental Design screening.
(D) Construction of the 3 screening experimental designs. Plackett-Burman designs were generated in 36,36 and 24 runs using R DOE package.

Supp Figure 2

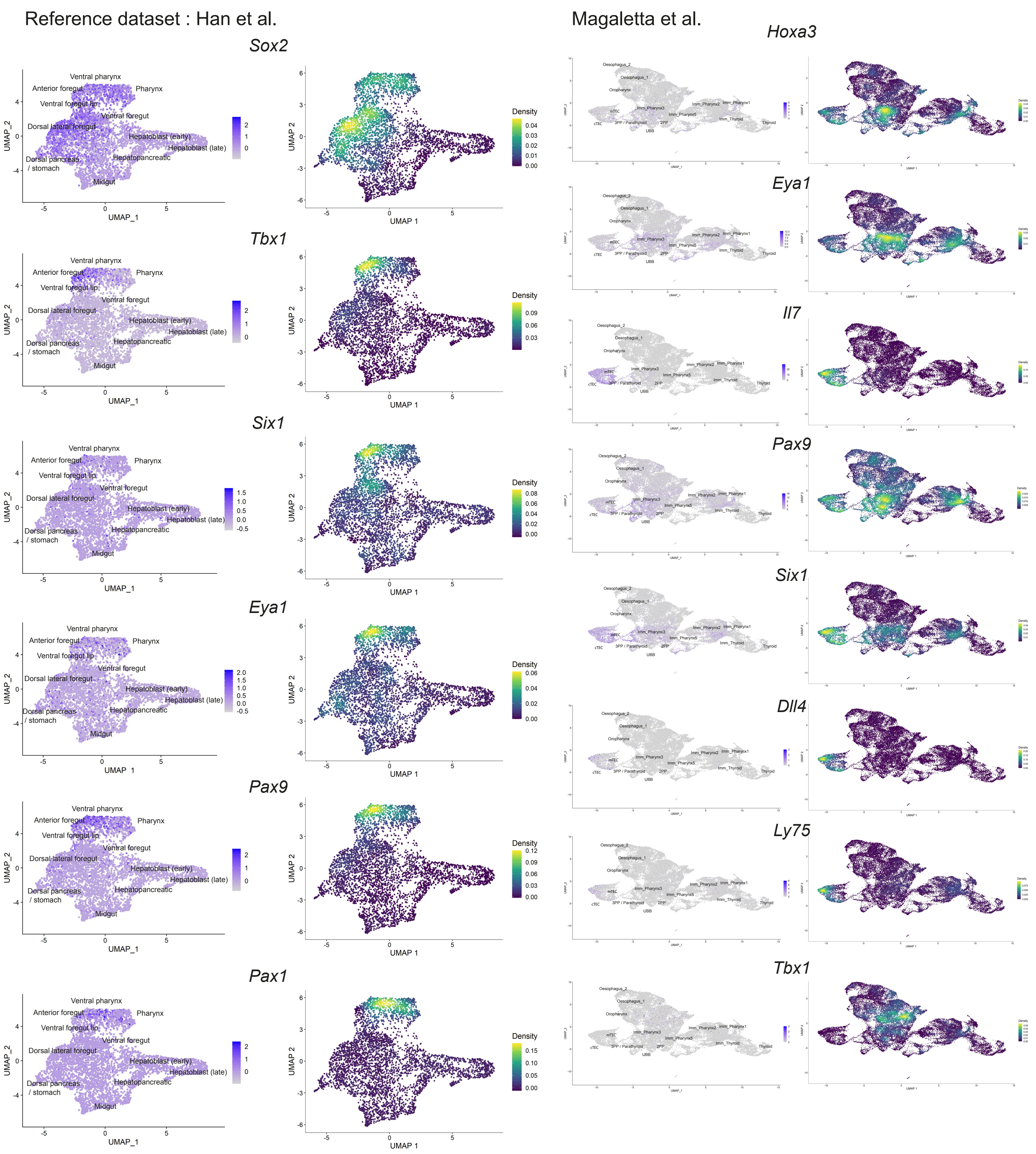


**Supplementary Figure 2.** Main markers expression in the Han et al. (*Left*) and Magaletta et al. (*Right*) datasets of pharyngeal organogenesis.

Supp Figure 3

Reference dataset : Han et al.

D5 → D7

| | Df | Sum Sq | Mean Sq | F value | Pr(>F) | |
|---|---|---|---|---|---|---|
| RA_J5 | 1 | 21.3842 | 21.3842 | 164.6310 | 4.345e-07 | *** |
| IWR1_J5 | 1 | 7.0510 | 7.0510 | 54.2835 | 4.246e-05 | *** |
| BMP4_J5 | 1 | 0.0149 | 0.0149 | 0.1149 | 0.74242 | |
| NOGGIN_J5 | 1 | 0.9652 | 0.9652 | 7.4308 | 0.02338 | * |
| LY3_J5 | 1 | 0.6292 | 0.6292 | 4.8438 | 0.05526 | . |

D7 → D11

| | Df | Sum Sq | Mean Sq | F value | Pr(>F) | |
|---|---|---|---|---|---|---|
| cyclo_J7 | 1 | 0.2184 | 0.2184 | 3.9790 | 0.1168165 | |
| CHIR99_J7 | 1 | 18.4014 | 18.4014 | 335.2933 | 5.233e-05 | *** |
| FLI06_J7 | 1 | 7.7451 | 7.7451 | 141.1231 | 0.0002875 | *** |
| IWR1_J7 | 1 | 3.1897 | 3.1897 | 58.1195 | 0.0015895 | ** |
| BMP4_J7 | 1 | 0.0198 | 0.0198 | 0.3605 | 0.5805726 | |
| FGF8_J7 | 1 | 0.0072 | 0.0072 | 0.1316 | 0.7351782 | |

Reference dataset : Magaletta et al.

D7 → D11

| | Df | Sum Sq | Mean Sq | F value | Pr(>F) | |
|---|---|---|---|---|---|---|
| cyclo_J7 | 1 | 0.0212 | 0.0212 | 0.8544 | 0.4076311 | |
| CHIR99_J7 | 1 | 13.9327 | 13.9327 | 562.1067 | 1.877e-05 | *** |
| FLI06_J7 | 1 | 13.9328 | 13.9328 | 562.1095 | 1.877e-05 | *** |
| IWR1_J7 | 1 | 3.3229 | 3.3229 | 134.0599 | 0.0003179 | *** |
| BMP4_J7 | 1 | 0.0509 | 0.0509 | 2.0538 | 0.2251125 | |
| FGF8_J7 | 1 | 0.0276 | 0.0276 | 1.1139 | 0.3507582 | |

D11 → D13

| | Df | Sum Sq | Mean Sq | F value | Pr(>F) | |
|---|---|---|---|---|---|---|
| EGF_J11 | 1 | 0.5055 | 0.5055 | 8.3921 | 0.101365 | |
| IGF1_J11 | 1 | 0.6645 | 0.6645 | 11.0324 | 0.079926 | . |
| RANKL_J11 | 1 | 0.6099 | 0.6099 | 10.1255 | 0.086185 | . |
| FGF10_J11 | 1 | 0.1955 | 0.1955 | 3.2455 | 0.213411 | |
| BMP4_J11 | 1 | 0.5809 | 0.5809 | 9.6437 | 0.089927 | . |

**Supplementary Figure 3.** DOE ANOVA results of the main indicidual effect of the tested factors

Supp Figure 4

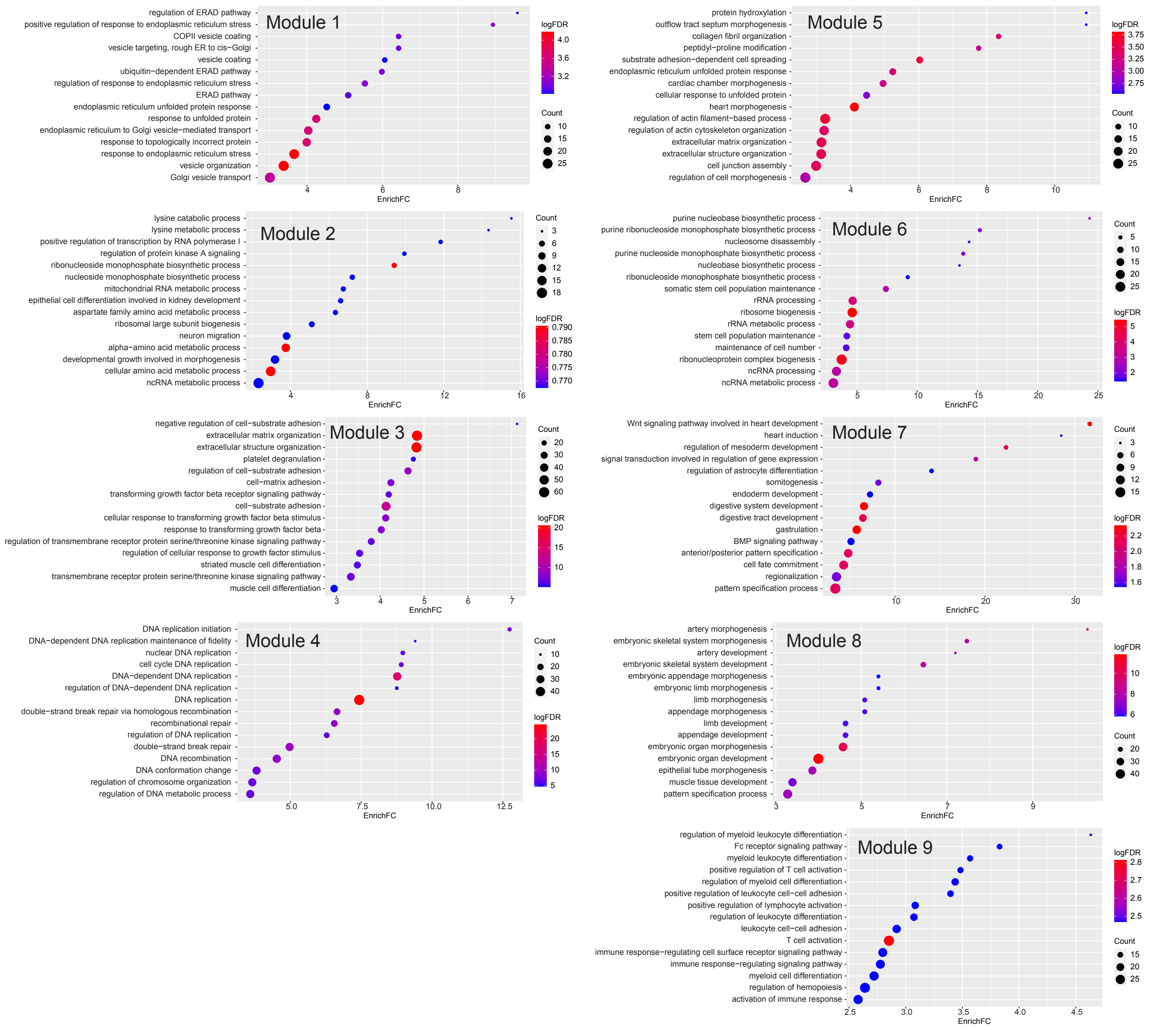


**Supplementary Figure 4.** Top 15 GO terms for Biological Processes enriched for the 9 differentially expressed gene modules from Fig 2.A

Supp Figure 5

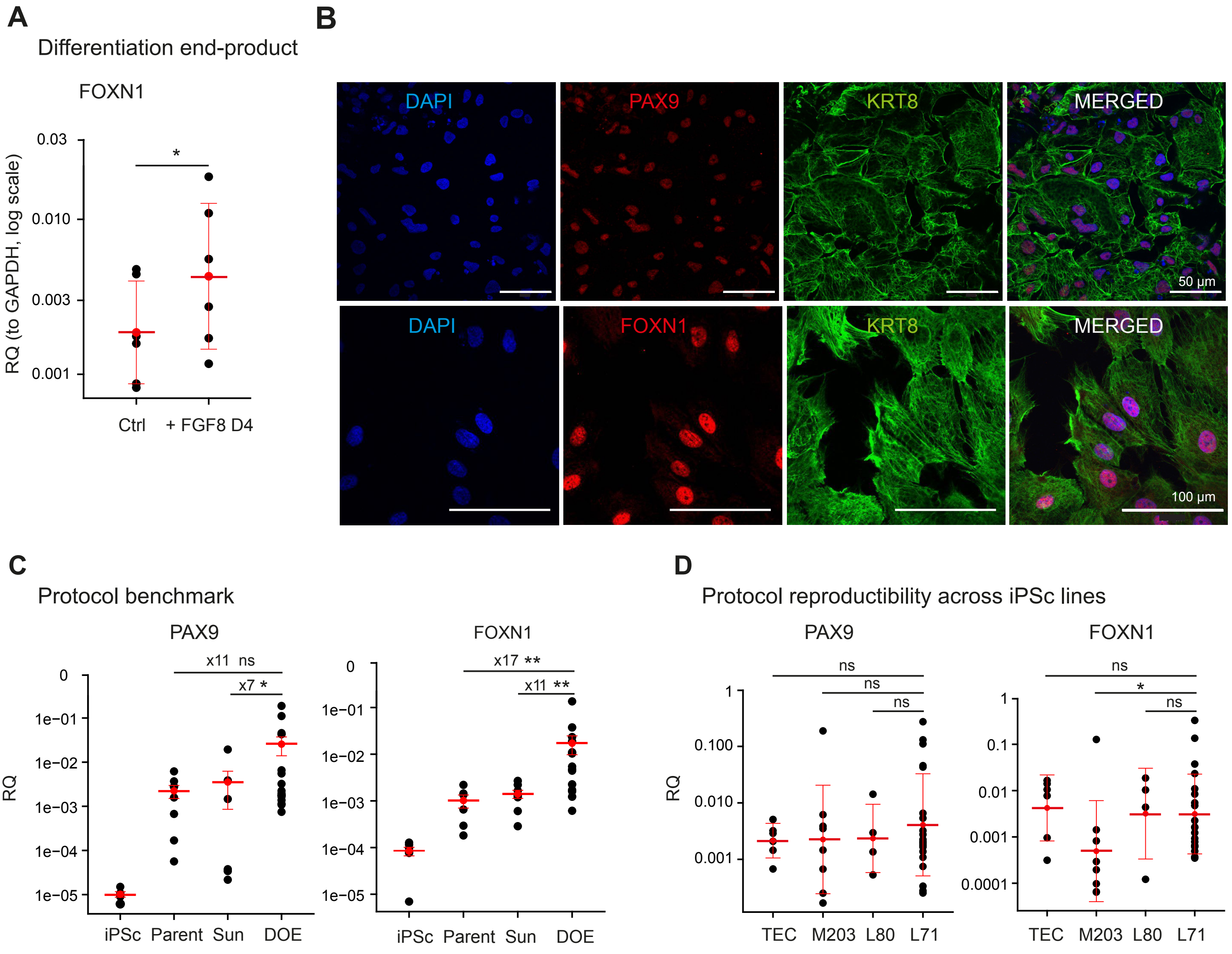


**E**

| Name | Specie | Cell type | Repro | Donor sex |
|---|---|---|---|---|
| hiN.Fm.m.LON71.019 | Human | Fibroblasts | ARNm | Male |
| hiN.Fm.f.LON80.002 | Human | Fibroblasts | ARNm | Female |
| hiN.Fs.f.MIPS203.B11 | Human | Fibroblasts | Sendai | Female |

**Suplementary Figure 5**. Validation of DOE transcriptomic result
(A) FGF8 supplementation at D4 improves FOXN1 expression in the differentiation product. Relative quantification by qPCR, using GAPDH expression ratio, (n = 6).
(B) Immunofluorescence imaging of thymic epithelium markers in TEP. Nuclei are stained with DAPI.
(C) qPCR quantification of TEP markers in our optimized protocol at D14 compared with two state-of-the-art protocols. Gene expression is measured as relative quantification of GAPDH, (n=6)
(D) Differentiation protocol is replicable to 3 different iPSc lines showing significant difference of expression by qPCR (relative quantification to GAPDH, log scale) only for FOXN1 in the MIPS203 line
(L80 : LON80, M203 : MIPS203, L71 : LON71, TEC : primary TEC control)
(E) iPSc cell line information

Supp Figure 6

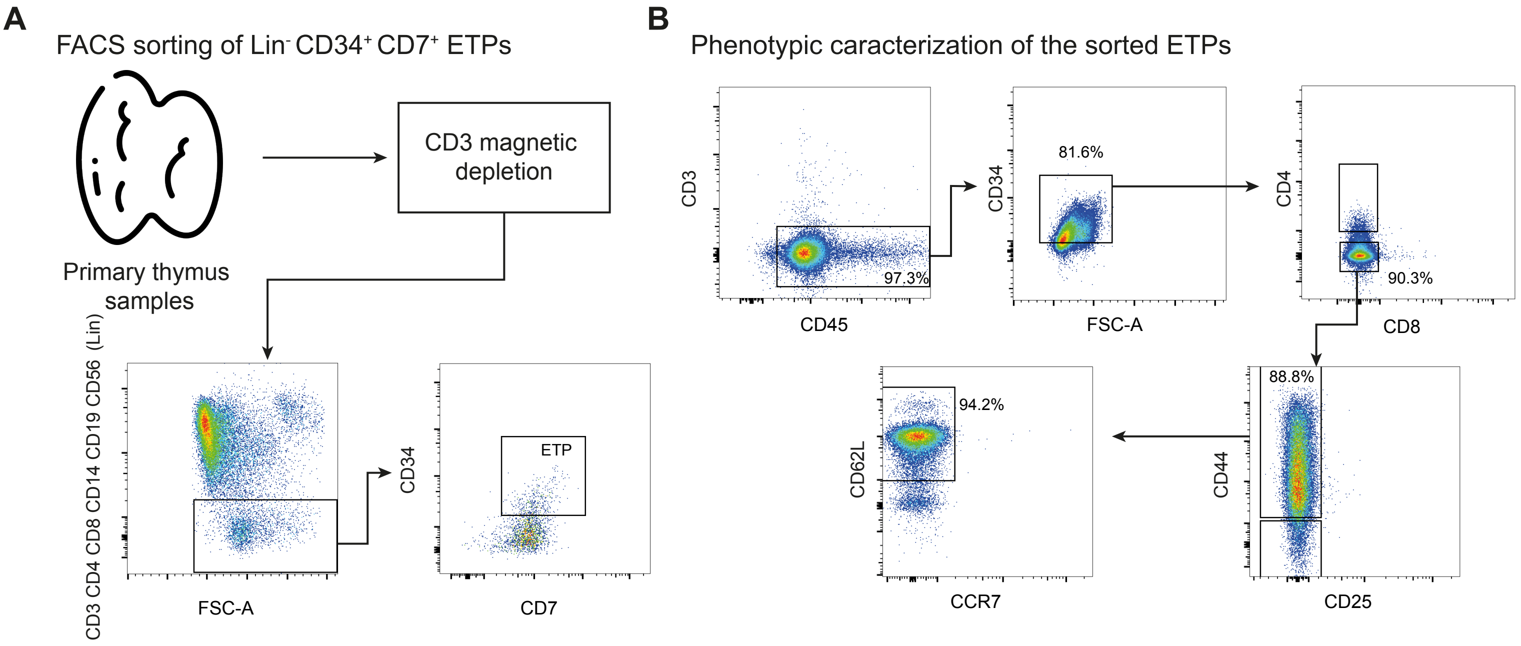


**Suplementary Figure 6**. Early thymocytes (ETP) isolation strategy.
(A) Sorting of human primary early thymic progenitors (ETP) characterized as CD3-CD4-CD8-CD14-CD19-CD56-(Lin)CD34+CD7+ by flow cytometry after enrichment by magnetic depletion of CD3+ thymocytes.
(B) FACS characterization of freshly sorted ETP show high purity of CD45+CD3-CD34+ ETP progenitors expressing CD44 and CD62L.

Supp Figure 7

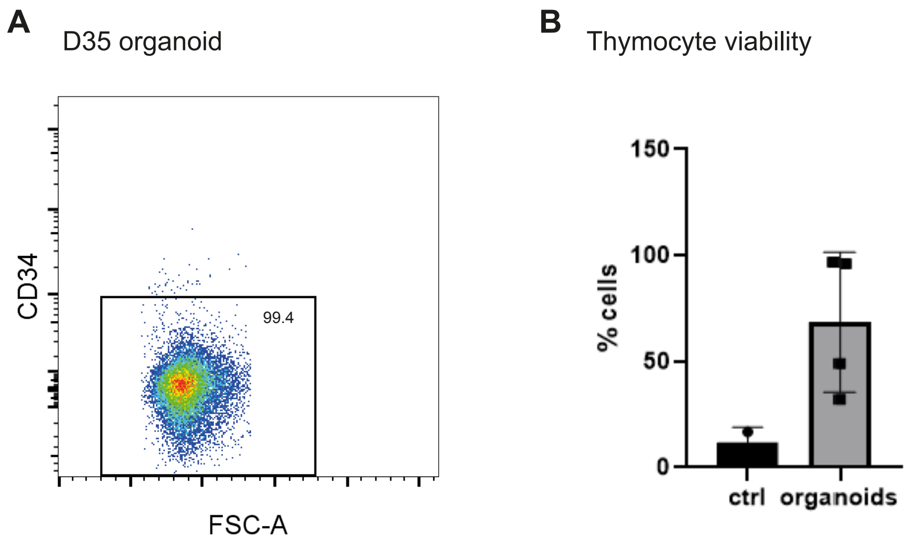


**Supplementary Figure 7.**
(A) Flow cytometry of the hematopoietic fraction of D35 hTO confirms loss of CD34
(B) Proportion of DAPI- cells in the hematopoetic product in hTOs

Supp Figure 8

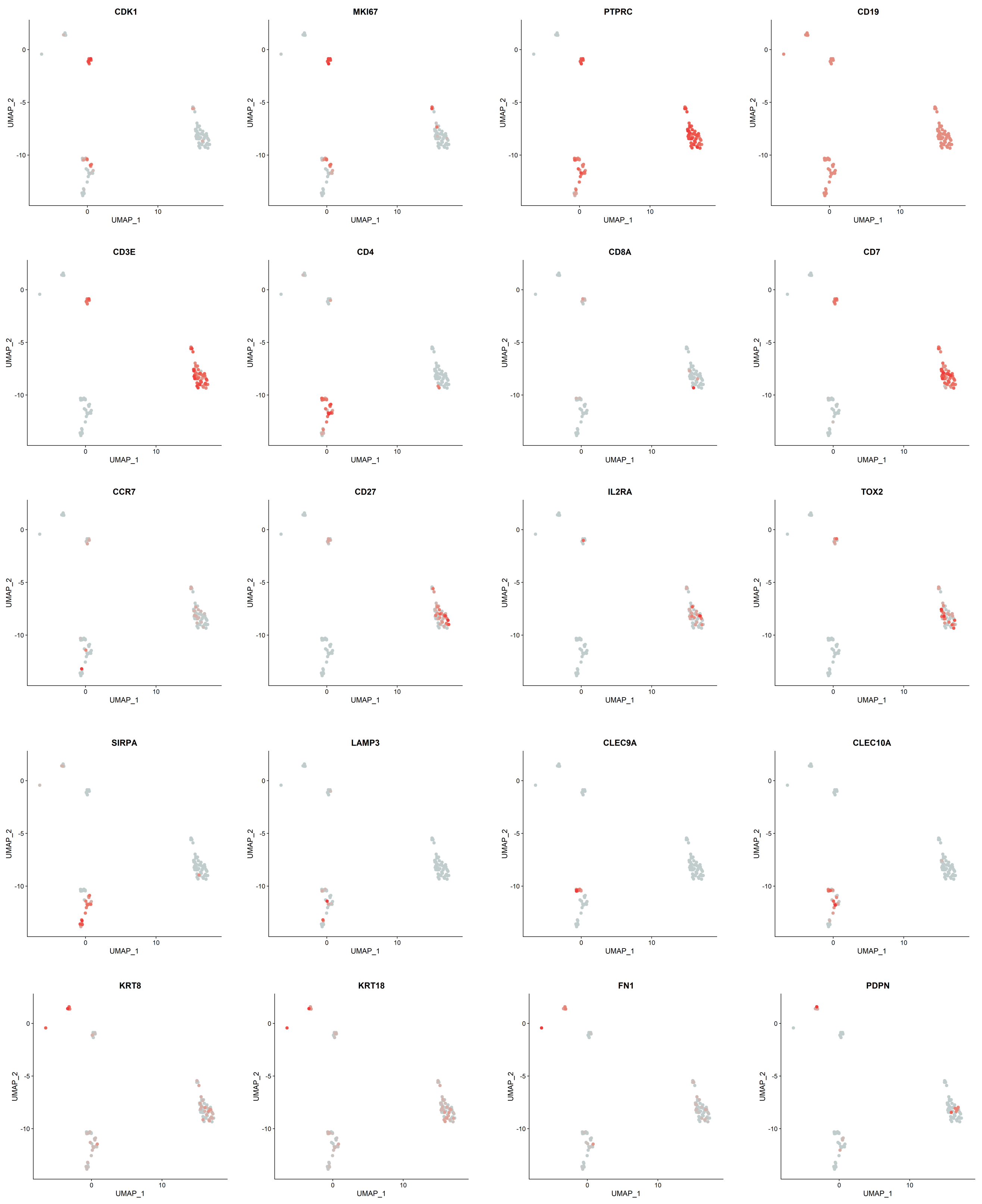


**Supplementary Figure 8 .** Main markers expression in our scRNA-seq data from a D28 hTO allows identification of a stromal cluster, two T cell clusters and one DC cluster.
